# Co-circulating mumps lineages at multiple geographic scales

**DOI:** 10.1101/343897

**Authors:** Shirlee Wohl, Hayden C. Metsky, Stephen F. Schaffner, Anne Piantadosi, Meagan Burns, Joseph A. Lewnard, Bridget Chak, Lydia A. Krasilnikova, Katherine J. Siddle, Christian B. Matranga, Bettina Bankamp, Scott Hennigan, Brandon Sabina, Elizabeth H. Byrne, Rebecca J. McNall, Daniel J. Park, Soheyla Gharib, Susan Fitzgerald, Paul Barriera, Stephen Fleming, Susan Lett, Paul A. Rota, Lawrence C. Madoff, Bronwyn L. MacInnis, Nathan L. Yozwiak, Sandra Smole, Yonatan H. Grad, Pardis C. Sabeti

**Affiliations:** Broad Institute of MIT and Harvard, Cambridge, MA, USA.; Center for Systems Biology, Department of Organismic and Evolutionary Biology, Harvard University, Cambridge, MA, USA.; Department of Electrical Engineering and Computer Science, Massachusetts Institute of Technology, Cambridge, MA, USA.; Department of Immunology and Infectious Diseases, Harvard T.H. Chan School of Public Health, Boston, MA, USA.; Division of Infectious Diseases, Department of Medicine, Massachusetts General Hospital, Boston, MA, USA.; Massachusetts Department of Public Health, Jamaica Plain, MA, USA.; Center for Communicable Disease Dynamics, Harvard T.H. Chan School of Public Health, Boston, MA, USA.; Division of Viral Diseases, Centers for Disease Control and Prevention, Atlanta, GA, USA.; Harvard University Health Services, Harvard University, Cambridge, MA, USA.; Department of Medicine, University of Massachusetts Medical School, Worcester, MA, USA.; Division of Infectious Diseases, Brigham and Women’s Hospital, Harvard Medical School, Boston, MA, USA.; Howard Hughes Medical Institute, Chevy Chase, MD, USA.

## Abstract

Despite widespread vaccination, eleven thousand mumps cases were reported in the United States (US) in 2016–17, including hundreds in Massachusetts, primarily in college settings. We generated 203 whole genome mumps virus (MuV) sequences from Massachusetts and 15 other states to understand the dynamics of mumps spread locally and nationally, as well as to search for variants potentially related to vaccination. We observed multiple MuV lineages circulating within Massachusetts during 2016–17, evidence for multiple introductions of the virus to the state, and extensive geographic movement of MuV within the US on short time scales. We found no evidence that variants arising during this outbreak contributed to vaccine escape. Combining epidemiological and genomic data, we observed multiple co-circulating clades within individual universities as well as spillover into the local community. Detailed data from one well-sampled university allowed us to estimate an effective reproductive number within that university significantly greater than one. We also used publicly available small hydrophobic (SH) gene sequences to estimate migration between world regions and to place this outbreak in a global context, but demonstrate that these short sequences, historically used for MuV genotyping, are inadequate for tracing detailed transmission. Our findings suggest continuous, often undetected, circulation of mumps both locally and nationally, and highlight the value of combining genomic and epidemiological data to track viral disease transmission at high resolution.

---

An unusually large number of mumps cases were reported in the US in 2016 and 2017, despite high rates of vaccination^1,2^. In the pre-vaccination era, mumps was a routine childhood disease, with over 180,000 cases reported in the US annually^1^. After the mumps vaccine was introduced in 1967, mumps incidence declined by more than 99%^1^. Case counts rose again briefly in the mid-1980s, and then continued to decrease after two Measles-Mumps-Rubella (MMR) vaccine doses were recommended in 1989 following a national outbreak of measles^3^. In the early 2000s, only a few hundred cases of mumps were observed annually in the US1. This low nationwide incidence was interrupted by a large outbreak (>5,000 cases) in the Midwestern US in 2006^4^, followed by a period of low incidence with minor outbreaks until 2016. The recent resurgence in mumps is not fully understood, although waning immunity^5^ and genetic changes in circulating viruses may be contributing factors.

Within Massachusetts, over 250 cases were reported in 2016 and more than 170 in 2017, far exceeding the usual state incidence of <10 cases per year^6^ (**Fig. 1a-b**). As seen in other recent outbreaks, most cases were associated with academic institutions and other close contact settings^4,7^. Mumps was reported in at least 18 colleges and universities in Massachusetts, including Harvard University (Harvard), University of Massachusetts Amherst (UMass), and Boston University (BU). Of the individuals infected, at least 65% had the recommended two doses of the MMR vaccine (**Extended Data Table 1a**).

**Fig. 1.**
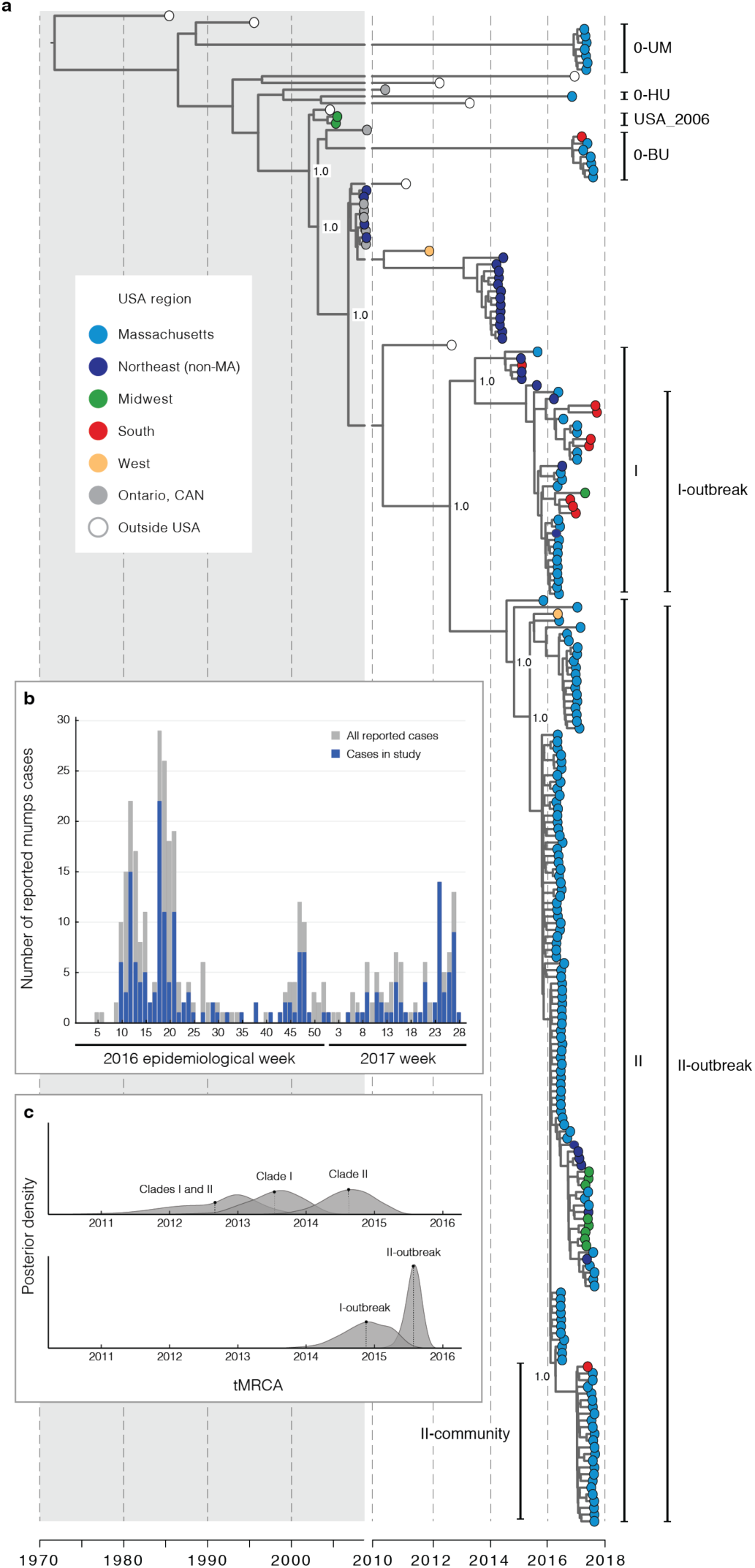
Mumps cases in Massachusetts and circulation of MuV in the United States. (**a**) Maximum clade credibility tree of 225 MuV genotype G whole genome sequences, including 200 generated in this study. Labels on selected internal nodes indicate posterior support. Clades I and II contain 91% of the samples from the 2016–2017 Massachusetts outbreak; I-outbreak and II-outbreak are the smallest clades within them that contain all samples from the outbreak. Clade 0-UM contains samples associated with UMass other than those in clades I and II; the same is true for 0-BU (BU) and 0-HU (Harvard). II-community is comprised primarily of samples associated with a local Massachusetts community. (**b**) Number of reported mumps cases by epidemiological week in Massachusetts (gray) and in this study (blue); samples included in this study are representative of all cases (see Extended Data Table 1). Samples from prior to 2016 are not shown. (**c**) Probability distributions for the tMRCA of selected sequences labeled in (a) (see **Extended Data Table 3** for additional clades). Dotted line: mean of each distribution.

We set out to investigate the evolution and spread of MuV, generating 203 whole genomes (160 collected in Massachusetts and 43 in 15 other states) through a combination of unbiased and capture-based sequencing approaches^8,9^ (see Methods, **Supplementary Table 1**). The genomes had a median of 99.48% unambiguous base calls and a median depth of 176x (**Extended Data Fig. 1a**). Technical replicates had high concordance: in 27 samples prepared more than once, only two base calls differed across replicates. We also sequenced a set of 29 PCR-negative samples from suspected mumps patients in Massachusetts to determine if we could detect MuV or other viruses that may explain their symptoms. We saw evidence for MuV in one sample, and identified 4 other viruses known to cause upper respiratory symptoms (at least two of which are known to cause parotitis) in four separate samples^10–12^ (**Extended Data Table 2**).

In light of the large number of vaccinated individuals who contracted MuV during this outbreak, we looked for genetic variation that could suggest vaccine escape (**Supplementary Table 2**). We focused on genotype G MuV genomes, the predominant genotype found in the US, and estimated d*N*/d*S* on 200 genotype G genomes from our dataset together with all 25 genotype G whole genomes available from NCBI GenBank^13^. This provided no strong evidence for positive selection at any specific site (**Extended Data Fig. 2a**, **Supplementary Table 3**) or in any gene (**Extended Data Fig. 2b**). No fixed nucleotide substitutions were associated with vaccine status or time since vaccination (**Extended Data Fig. 2c,e**). We also investigated changes in immunogenic regions of the mumps genome between our samples and the vaccine strain used in the US, focusing on the hemagglutinin (HN) protein, the primary target of neutralizing antibodies^14^. We found numerous fixed differences (**Extended Data Fig. 2d**), including in regions of potential immunological significance, but did not find clear evidence for variants that might lead to escape from vaccine-induced immunity (see **Supplementary Information**).

We investigated mumps transmission at multiple geographic scales, first examining how the Massachusetts outbreak fits into the larger context of resurgent mumps. We performed a phylogenetic analysis using the same 225 genotype G MuV genomes as above (**Extended Data Fig. 3a**). The resulting phylogeny (**Fig. 1a**) suggests that most MuV genomes from cases in Massachusetts descend from those in the 2006 US outbreak: these Massachusetts genomes fall within a clade that contains the 2006 samples, and the root of this clade lies within or very close to the 2006 samples. This conclusion is supported by a root-to-tip analysis (**Extended Data Fig. 3b,c**). Other US samples collected between 2006 and the Massachusetts outbreak (2016) also fall within this clade, suggesting sustained MuV transmission within the country. Additionally, we saw that MuV sequences from Massachusetts are interspersed with sequences across the US also collected in 2016–2017 (**Fig. 1a**), providing evidence for extensive geographic movement of MuV within the country on short time scales. A principal components analysis of MuV sequences (**Extended Data Fig. 3d**) similarly shows that sequences from other regions within the US often cluster with ones from Massachusetts.

We see clear evidence for several introductions of MuV into Massachusetts, including two distinct MuV lineages (clades I and II) that descend from the 2006 outbreak and diverged late in 2012 (**Fig. 1c, Extended Data Table 3**). To estimate when the two primary clades entered Massachusetts, we calculated the time to the most recent common ancestor (tMRCA) of each using only samples from the 2016–2017 Massachusetts outbreak (I-outbreak = July 2015, II-outbreak = November 2014) (**Fig. 1c, Extended Data Table 3**). These tMRCAs, as well as the estimated divergence time of the two clades, are well before the diagnosis of the first cases associated with the 2016–2017 outbreak, providing strong evidence for a minimum of two introductions into Massachusetts. We also saw evidence for additional introductions: three clades (0-UM, 0-HU, 0-BU) contain MuV sequences distinct from the majority of Massachusetts sequences, and each has at least one MuV sequence from a patient with foreign travel history during the incubation period of the virus (12–25 days before symptom onset^15^).

The phylogeny illuminates details of the outbreak at finer scales as well, showing that multiple clades were co-circulating within academic institutions. Specifically, there are multiple viral lineages in individuals associated with UMass (clade II and 0-UM), in Harvard-associated individuals (clades I, II, and 0-HU), and in BU-associated individuals (clades I, II, and 0-BU) (**Extended Data Fig. 4a**). Thus, what appeared to be a single outbreak across multiple institutions is shown by sequence data to consist of multiple overlapping chains of transmission. This adds to the overall picture of mumps in the US as being sustained by multiple, widespread lineages that lead to local flare-ups.

Viral genome sequence data allowed us to identify a spillover event from one institution into a non-university community. Before genomic data were available, cases associated with Harvard and with an apparently unrelated local community (clade II-community) were inferred to be separate outbreaks due to the different populations affected (mostly students versus adults with no obvious university connection) and an apparent five month gap between the two sets of cases (**Supplementary Table 1**). From the phylogeny, however, it is clear that these two groups of cases are connected, and that the community-associated cases likely represent a spillover from Harvard into the broader population (**Fig. 2a**). Additional epidemiological investigation identified three infected individuals associated with both Harvard and the community who could have served as transmission links. The gap in time between these two sets of linked cases suggests local undetected mumps circulation, potentially due to unreported cases or asymptomatic infection^16^. In line with the picture seen above, this observation illustrates sustained transmission that is only sporadically detected.

**Fig. 2.**
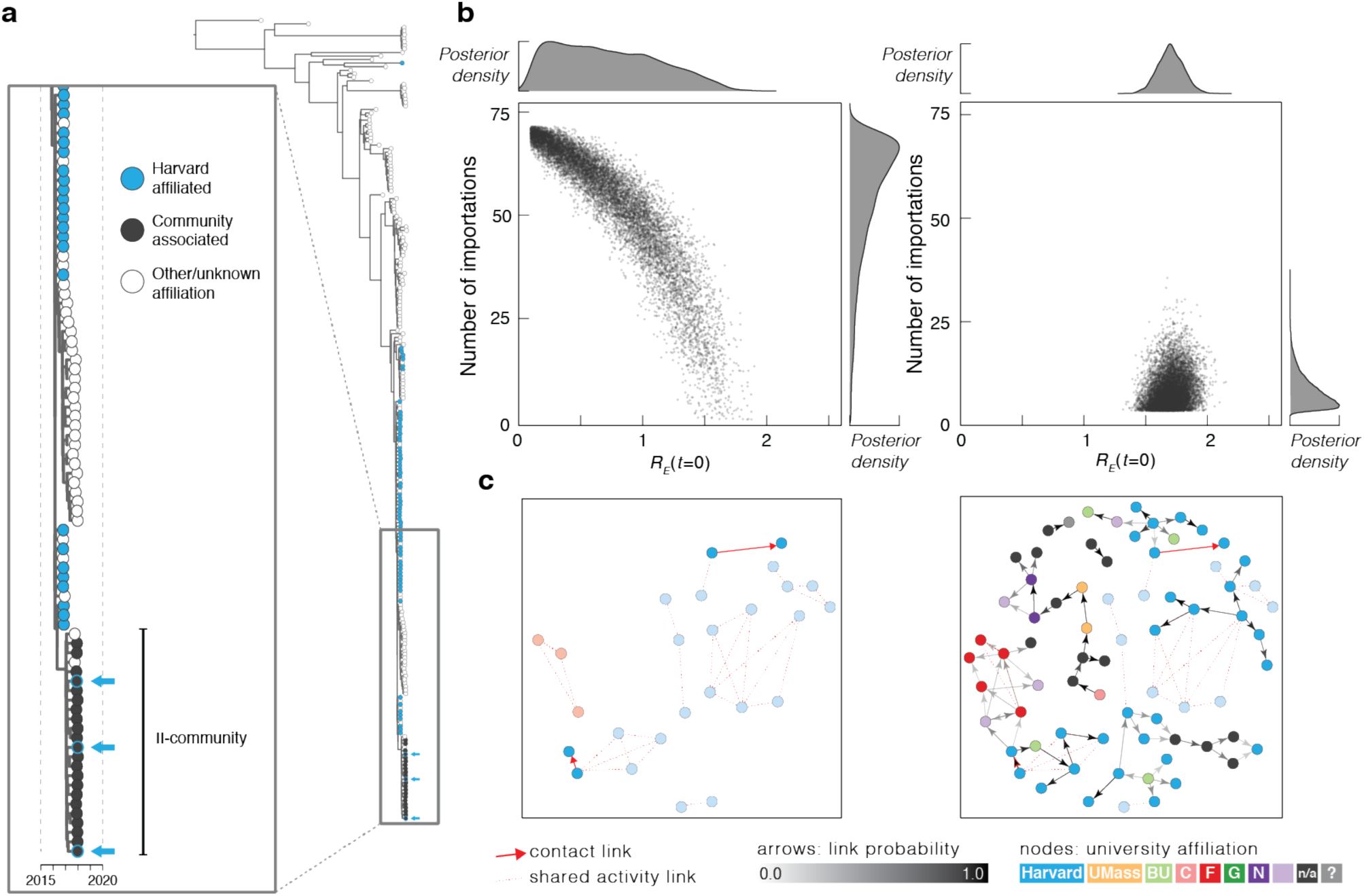
Epidemiological modeling and transmission reconstruction. (**a**) Zoom view of the clade II-community and its ancestors (see **Fig. 1a**), colored by academic institution affiliation. Arrows highlight samples from 3 individuals affiliated with both the II-community and Harvard. (**b**) Number of importations into Harvard calculated without (left) and with (right) viral genetic information as input. Each point represents a sample from the posterior distribution of *R_E_*(*t=*0) and the number of introductions, based on simulated transmission dynamics. (**c**) Transmission reconstruction of individuals within clade II-outbreak; samples are colored by institution affiliation (light purple: other institution; n/a: no affiliation; question mark: unknown affiliation). Left: reconstruction using epidemiological data only; all individuals in clade II-outbreak with known epidemiological links (red arrows) are shown. Right: reconstruction using MuV genomes and collection dates. Arrow shading indicates probability of direct transmission between individuals (minimum probability shown: 0.3); individuals with no estimated links are not shown. Arrows outlined in red represent transmission events identified by both genomic and epidemiological data. Faded nodes are those only connected by shared activity links (i.e. no inferred or known direct transmission).

The comprehensive sampling of clinical mumps cases at Harvard again allowed us to combine genomic and epidemiological data to better understand outbreak dynamics, this time by estimating the number of MuV introductions into the university. The detailed epidemiological data alone were inadequate to distinguish whether the initial effective reproductive number, *R_E_*(*t*=0), within the university exceeded the critical threshold of 1.0 (**Fig. 2b left, Extended Data Fig. 5**). However, incorporating the observed number of distinct viral lineages observed markedly improved our ability to infer transmission dynamics within the university, supporting an estimate of five (95% CI: 4–18) distinct introductions, each expected to cause *R_E_*(*t*=0)=1.70 (95% CI: 1.50–1.91) secondary infections (**Fig. 2b right**). That *R_E_*(*t*=0) is well above 1.0 has implications for the required reach of outbreak response vaccination.

We next investigated the usefulness of sequence data to supplement epidemiological data at the finest analysis scale: reconstructing individual transmission chains. Epidemiological links alone paint a sparse picture of transmission (**Fig. 2c left**). To determine how effective genomic data could be in inferring any missing epidemiological links, we examined the genetic distance between samples with a known epidemiological link (i.e., samples likely part of the same transmission chain). Samples with a known link group together according to genetic distance (**Extended Data Fig. 6a**), and genetic distance can be used as a predictor of epidemiological linkage (**Extended Data Fig. 6b**). Given this, we used genomic data (along with sampling dates, but without using known epidemiological links) to reconstruct transmission chains during the outbreak (**Fig. 2c right**), focusing on samples within clade II-outbreak. These reconstructed transmission chains in several cases support known epidemiological links, but also included links between individuals without known contacts. The reconstruction did not reproduce several non-direct contact epidemiological links (‘shared activity links’, see Methods), which highlights the utility of both epidemiological and genetic data in an outbreak context.

When shared between samples, within-host variants (intrahost variants, or iSNVs) can provide additional information about transmission chains during viral outbreaks^17–19^, but proved to be uninformative in our dataset. We found no evidence for transmitted iSNVs among our samples, including among sequences from five direct contacts (**Supplementary Table 2**). These data suggest that the MuV transmission bottleneck may be small enough to preclude shared within-host variation (see **Supplementary Information**). We did, however, detect changes to MuV during the course of individual infections. In the two patients for whom we had multiple samples, the MuV genomes differed at one site between time points (in both cases, nine days apart) (see **Supplementary Information**).

To understand MuV circulation in its global context, we analyzed the 316-nucleotide SH gene. This gene was originally chosen for MuV genotyping^20,21^ because it has a high density of variation and provides a small, convenient target amenable to sequencing methods available at the time; it is therefore the region of the genome for which the most sequence data is available. Our SH dataset consists of 3,646 SH sequences available on NCBI GenBank from around the world, including those from genomes generated in this study (**Fig. 3a, Extended Data Fig. 7a**). We calculated tMRCA for each of the 11 clinically relevant genotypes (**Fig. 3b**) and found that genotype A, the strain used in most vaccines (including the Jeryl Lynn vaccine used in the US), coalesces the earliest and appears to have stopped circulating (previously noted in ref. 22). The other genotypes, including genotype G, coalesce over a decade later.

**Fig. 3.**
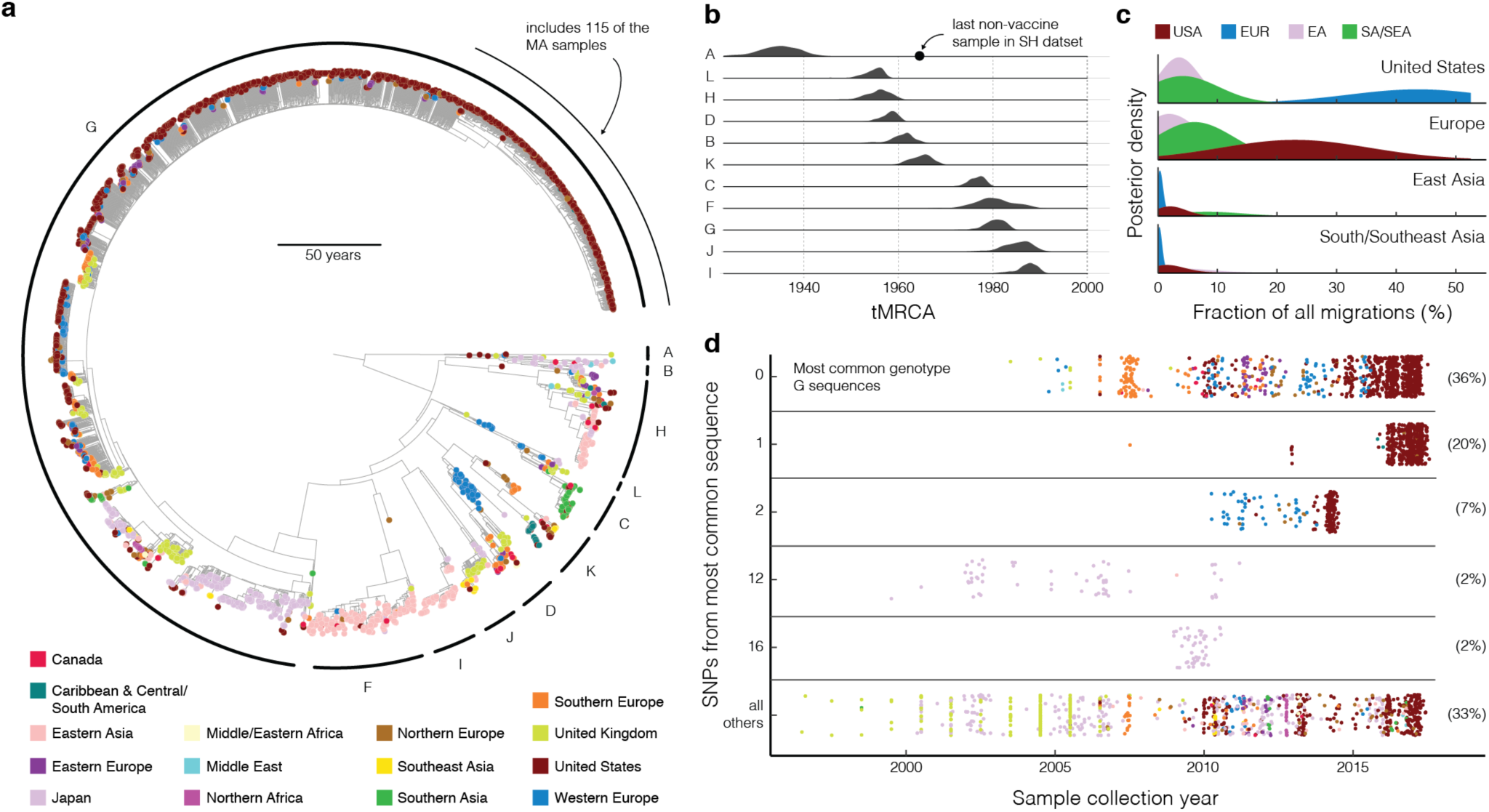
Global spread of MuV based on SH gene sequences. (**a**) Maximum clade credibility tree of 3,646 publicly available SH gene sequences, including 193 generated in this study. (**b**) Probability distribution for the tMRCA of each included genotype. The date of the most recent genotype A clinical sample is indicated, excluding samples that closely resemble a MuV vaccine strain. (**c**) Migration between 4 global regions. Each plot shows a posterior probability density, taken across resampled input, of the fraction of all reconstructed migrations that occur to the destination (indicated in upper right) from each of the other 3 sources. (**d**) Identical genotype G sequences over time (two samples collected before 1995 are not shown). Each dot represents a sample (colored as in (a)) and each row contains samples with identical SH sequences, except the bottom, which includes samples whose sequences are distinct from those in the above 5 categories. Numbers right of each row are the percentage of all genotype G samples found in that row.

We performed a phylogeographic analysis on these SH sequences to understand movement of MuV between world regions. To reduce temporal and geographic sampling biases due in part to incomplete global MuV surveillance, we looked at samples collected since 2010 in four well-sampled regions (US, Europe, East Asia, and South/Southeast Asia). (For details regarding resampling, see Methods.) We show significant MuV migration between the US and Europe (**Fig. 3c, Extended Data Fig. 7b,c**) and see that most introductions to the US are from Europe, although there is also support for mumps migrations from East Asia and South/Southeast Asia to the US. Likewise, most introductions to Europe appear to be from the US. We also found that recent sequences from the US have primarily European ancestry, and vice versa (**Extended Data Fig. 7d**). Notably, recent sequences from East Asia and South/Southeast Asia have minimal external ancestry, indicating relatively little spread to these regions (**Fig. 3c**).

It is important to note that the SH gene is less than 3% of the MuV genome, represents only a fraction of MuV genetic diversity, and has fewer phylogenetically relevant nucleotide substitutions than other genes^23^. While these sequences are useful for tracking genotype spread on a global scale, the lack of variability across them (**Fig. 3d**) may inflate our estimates of migration. Moreover, lack of variability makes it difficult or impossible to reconstruct MuV spread on smaller temporal and geographic scales. For example, analysis using SH gene sequencing alone would not have shown a genetic link between Harvard and the geographic community (**Extended Data Fig. 6c, Extended Data Fig. 8**). Furthermore, there is limited variability in genotype G sequences in the US, even over a time period of several years (**Fig. 3d**), highlighting the value of whole genome sequencing.

The pairing of whole genome sequences with epidemiological data provides a detailed picture of outbreak dynamics at a full range of geographic scales, from national migration patterns to individual transmission chains. Our MuV sequence data show that multiple co-circulating strains underlie recent increases in mumps cases within Massachusetts, and even within single institutions. Unexpected insights using genomic epidemiology include the finding that two apparently unrelated outbreaks were in fact linked and that a sustained series of cases within one institution stemmed from multiple independent introductions. While SH gene sequencing has proved useful for tracking broad trends in MuV circulation, many of our findings — continuous circulation of MuV lineages, connections between cases in Massachusetts, specific dynamics of MuV at Harvard — were only possible through the use of whole genome sequencing (**Extended Data Fig. 8**).

The large number of mumps cases among vaccinated individuals has raised concerns about mumps vaccination. The US Advisory Committee on Immunization Practices recently recommended a third dose of MMR vaccine for persons at increased risk for acquiring mumps because of an outbreak^24^. Our finding of ongoing, undetected MuV circulation in the US gives additional weight to this recommendation, and underscores the importance of interrupting transmission in high-risk communities. This study presents a model for the future of genomics-informed infectious disease control, in which genomic data can both support and extend conventional epidemiological data to inform our understanding of how pathogens spread.

## Acknowledgements

We thank J. Qu and R. Shah for sample preparation support; A. Matthews and S. Winnicki for management and guidance; I. Shlyakhter, S. Weingarten-Gabbay, S. Ye, C. Tomkins-Tinch, and the Sabeti Laboratory for discussions and reading of the manuscript; J. Hall, P. Patel, E. Buzby, K. Chen, and F. Halpern-Smith for mumps diagnosis and laboratory support; A. Osinski, C. Brandeburg, H. Johnson, J. Cohen, K. Royce, M. Popstefanija, N. Harrington, R. Hernandez, and J. Leaf for case management and epidemiological investigation; T. Mason and the Broad Institute Genomics Platform for sequencing support; M. Salit for sharing reagents. We are indebted to mumps patients and clinical and epidemiological teams for making this work possible.

Funding was provided by: NIH NIAID U19AI110818 (Broad Institute); NIH NIAID U54GM088558 (J.A.L.); Howard Hughes Medical Institute (P.C. S.); Harvard University Burke Global Health Fellowship (P.C.S.); Amazon Web Services Cloud Credits for Research (P.C.S.). The project described was supported by award number T32GM007753 from the National Institute of General Medical Sciences (E.H.B.). The content is solely the responsibility of the authors and does not necessarily represent the official views of the National Institute of General Medical Sciences or the National Institutes of Health.

## Author Contributions

S.W., A.P., K.J.S., C.B.M., B.B., S.H., and B.S. performed laboratory experiments and prepared samples for sequencing. H.C.M., K.J.S., and C.B.M. developed methods for MuV detection, targeted enrichment, and/or sequencing library preparation. S.W., H.C.M., S.F.S., A.P., M.B., J.A.L., B.C., L.A.K., K.J.S., and E.H.B. performed sequence assembly, curation, and/or data analyses. M.B., R.J.M., S.G., S.Fi., P.B., S.Fl., S.L., P.A.R., L.M., and S.S. oversaw clinical data and sample collection, and/or performed epidemiological investigation. J.A.L. developed the transmission model. D.J.P., S.G., S.Fi., P.B., S.Fl., S.L., P.A.R., L.M., B.L.M., N.L.Y., S.S., Y.H.G., and P.C.S. provided critical insights and guidance. S.W., B.C., P.A.R., L.M., B.L.M., N.L.Y., S.S., Y.H.G., and P.C.S oversaw study design and management. S.W., H.C.M., S.F.S., A.P., J.A.L., L.A.K., B.L.M., N.L.Y., Y.H.G., and P.C.S. drafted the manuscript. All authors reviewed the manuscript.

## Methods

### Data availability

The 203 MuV whole genome sequences generated in this study, as well as nine low quality sequences not included in the analysis, are available on NCBI GenBank^13^ under BioProject accession PRJNA394142 (accession numbers MF965196–MF965318 and MG986380–MG986468).

### Ethics statement

The study protocol was approved by the Massachusetts Department of Public Health (MDPH), Centers for Disease Control and Prevention (CDC), and Massachusetts Institute of Technology (MIT) Institutional Review Boards (IRB). Harvard University Faculty of Arts and Sciences and the Broad Institute ceded review of sequencing and secondary analysis to the MDPH IRB through authorization agreements. The MDPH IRB waived informed consent given this research met the requirements pursuant to 45 CFR 46.116 (d). The CDC IRB determined this project to be non-human subjects research as only de-identified leftover diagnostic samples were utilized. In compliance with the IRB agreement, Harvard University, University of Massachusetts Amherst, and Boston University granted approval for publication of their institution names in this paper.

### Sample collections and study subjects

Buccal swab samples were obtained from suspected and confirmed mumps cases tested at MDPH and CDC. Samples from MDPH (‘Cases in Study’, **Fig. 1b**) include all cases with a positive MuV PCR result (see ‘PCR diagnostic assays performed at MDPH and CDC’ below) collected between 1 January 2014 and 30 June 2017. Demographic information for all cases reported in Massachusetts (**Fig. 1b, Extended Data Table 1**) includes all confirmed and probable mumps cases reported to MDPH in that time period. Probable cases include cases with a positive mumps IgM assay result, or those with an epidemiological link to a confirmed case^25^. Samples from CDC are a selection of PCR-positive cases submitted to the CDC for testing between 2014–2017. See **Supplementary Table 1** for de-identified information, including metadata, about study participants.

### Viral RNA isolation

Sample inactivation and RNA extraction were performed at the MDPH, Broad Institute, and CDC. At MDPH, viral samples were inactivated by adding 300 µL Lysis/Binding Buffer (Roche) to 200 µL sample, vortexing for 15 seconds, and incubating lysate at room temperature for 30 minutes. RNA was then extracted following the standard external lysis extraction protocol from the MagNA Pure LC Total Nucleic Acid Isolation Kit (Roche) using a final elution volume of 60 µL. At the Broad Institute, samples were inactivated by adding 252 µL Lysis/Binding Buffer (ThermoFisher) to 100 µL sample. RNA was then extracted following the standard protocol from the MagMAX Pathogen RNA/DNA Kit (ThermoFisher) using a final elution volume of 75 µL. At CDC, RNA extraction followed the standard protocol from the QiaAmp Viral RNA mini kit (Qiagen).

### PCR diagnostic assays performed at MDPH and CDC

Diagnostic tests for presence of MuV were performed at the MDPH and CDC using the CDC Real-Time (TaqMan) RT-PCR Assay for the Detection of Mumps Virus RNA in Clinical Samples^7,26^. Each sample was run in triplicate using both the Mumps N Gene assay (MuN) and RNase P (RP) assay using this protocol. RT-PCR was performed on the Applied Biosystems 7500 Fast Real-Time PCR system or Applied Biosystems Prism 7900HT Sequence Detection System instrument.

### PCR quantification assays performed at Broad Institute

MuV RNA was quantified at the Broad Institute using the Power SYBR Green RNA-to-Ct 1-Step qRT-PCR assay (Life Technologies) and CDC MuN primers. The 10 µL assay mix included 3 µL RNA, 0.3 µL each MuV forward and reverse primers at 5 µM concentration, 5 µL 2x Power SYBR RT-PCR Mix, and 0.08 µL 125x RT Enzyme Mix. The cycling conditions were 48C for 30 min and 95C for 10 min, followed by 45 cycles of 95C for 15 sec and 60C for 30 sec with a melt curve of 95C for 15 sec, 55C for 15 sec, and 95C for 15 sec. RT-PCR was performed on the ThermoFischer QuantStudio 6 instrument. To determine viral copy number, a double-stranded gene fragment (IDT gBlock) was used as a standard. This standard is a 171 bp fragment of the MuV genome (GenBank accession: NC_002200) including the amplicon (sequence: GGA TCG ATG CTA CAG TGT ACT AAT CCA GGC TTG GGT GAT GGT CTG TAA ATG TAT GAC AGC GTA CGA CCA ACC TGC TGG ATC TGC TGA TCG GCG ATT TGC GAA ATA CCA GCA GCA AGG TCG CCT GGA AGC AAG ATA CAT GCT GCA GCC AGA AGC CCA AAG GTT GAT TCA AAC).

23S rRNA content in samples was quantified using the same Power SYBR Green RNA-to-Ct 1-Step qRT-PCR assay kit and cycling conditions. Primers were used to amplify a 183 bp universally conserved region of the 23S rRNA (fwd: 93a - GGG TTC AGA ACG TCG TGA GA, rev: 97ar - CCC GCT TAG ATG CTT TCA GC)^27^. To determine viral copy number, a double-stranded gene fragment (IDT gBlock) was used as a standard. This standard is a 214 bp fragment of the Streptococcus HTS2 genome (accession: NZ_CP016953) (sequence: AGC GGC ACG CGA GCT GGG TTC AGA ACG TCG TGA GAC AGT TCG GTC CCT ATC CGT CGC GGG CGT AGG AAA TTT GAG AGG ATC TGC TCC TAG TAC GAG AGG ACC AGA GTG GAC TTA CCG CTG GTG TAC CAG TTG TCT CGC CAG AGG CAT CGC TGG GTA GCT ATG TAG GGA AGG GAT AAA CGC TGA AAG CAT CTA AGT GTG AAA CCC ACC TCA AGA T). qPCR data from both assays — each performed only on a subset of samples — is reported in **Supplementary Table 1**.

### Bacterial rRNA depletion

Bacterial rRNA was depleted from some RNA samples using the Ribo-Zero Bacteria Kit (Illumina). At the hybridization step, the 40 µL reaction mix included 5 µL RNA sample, 4 µL Ribo-Zero Reaction Buffer, 8 µL Ribo-Zero Removal Solution, 22.5 µL water, and 0.5 µL synthetic RNA (25 fg) used to track potential cross-contamination (gift from M. Salit, NIST). Bacterial rRNA-depleted samples were purified using 1.8x volumes Agencourt RNAClean XP beads (Beckman Coulter) and eluted in 10 µL water for cDNA synthesis.

### Illumina library construction and sequencing

cDNA synthesis was performed as described in previously published RNA-seq methods^8^. In samples where bacterial rRNA was not depleted, 25 fg synthetic RNA was added at the beginning of cDNA synthesis to track sample cross-contamination. Positive control libraries were prepared from a mock MuV sample in which cultured Enders strain (ATCC VR-106) MuV was spiked into a composite buccal swab sample from healthy patients and diluted to MuV RT-qPCR Ct=21. This mock sample was extracted using the viral RNA isolation protocol described above, except that total nucleic acid was eluted in 100 µL. Negative control libraries were prepared from water. Illumina Nextera XT was used for library preparation: indexed libraries were generated using 16 cycles of PCR, and each sample was indexed with a unique barcode. Libraries were pooled equally based on molar concentration and sequenced on the Illumina HiSeq 2500 (100 or 150 bp paired-end reads) platform.

### Hybrid capture

Viral hybrid capture was performed as previously described^8^ using two different probe sets. In one case, probes were created to target MuV and measles virus (V-MM probe set), and in one case, probes were created to target 356 species of viruses known to infect humans (V-All probe set)^9^. Capture using V-All was used to enrich viral sequences primarily in samples in which we could not detect MuV, as well as in other samples (see **Supplementary Table 1** for a list of which samples were captured using which probe set). As described in ref. 9, the probe sets were designed to capture the diversity across all publicly available genome sequences on GenBank for these viruses. Probe sequences can be downloaded here: https://github.com/broadinstitute/catch/tree/cf500c69/probe-designs.

### Genome assembly

We used viral-ngs v1.18.1^28^ to assemble genomes from all sequencing runs. We used a set of MuV sequences (accessions: JX287389.1,FJ211586.1, AB000386.1, JF727652.1, AY685920.1,AB470486.1, GU980052.1, NC_002200.1, AF314558.1, AB823535.1, AF467767.2) to taxonomically filter these reads. We *de novo* assembled reads and scaffolded against the MuV genome with accession JX287389.1 to assemble a genome for each replicate. Then, we pooled read data from all sequencing replicates of each sample and repeated this assembly process to obtain final genomes. Each time we ran viral-ngs, we set the ‘assembly_min_length_fraction_of_reference’ and ‘assembly_min_unambig’ parameters to 0.01.

We replaced deletions in the coding regions with ambiguity (‘N’). In one sample, MuVs/Massachusetts.USA/11.16/5[G], with an insertion at position 3,903 (based on a full 15,384-nucleotide MuV genome, e.g., accession JN012242.1) we removed a poorly-supported (<5 reads covering the site) extra ‘A’ in a homopolymer region.

To calculate sequencing metrics (**Extended Data Fig. 1**), we used SAMtools^29^ to downsample raw reads for each replicate to 1 million reads and then re-ran assembly as described above. Samples from one contaminated sequencing batch were excluded, as were all replicates from PCR-negative samples. In cases where samples from two time points were sequenced from a single patient, we included only the first time point in the collection collection interval analysis (**Extended Data Fig. 1d**).

### Metagenomic analysis

We used the V-All probe set for capture on all samples from suspected mumps cases with a negative MuV PCR result (*n*=29). A subset of PCR-positive samples were also sequenced with this probe set (*n*=145) or without capture (‘unbiased’, *n*=111). We used the mock Enders strain MuV sample as a positive control on a sequencing run containing all PCR-negative samples, as well as a water sample as a negative control. We used the metagenomic tool Kraken v0.10.6^30^ via viral-ngs to identify the presence of viral taxa in each sample. We built a database similar to the one described in ref. ^9^, except without insect species. This database encompasses the known diversity of viruses known to infect humans and can be downloaded in three parts at https://storage.googleapis.com/sabeti-public/meta_dbs/kraken_full_20170522/[*file*] where [*file*] is: database.idx.lz4 (595 MB), database.kdb.lz4 (75 GB), and taxonomy.tar.lz4 (66 MB). Due to the possibility of contamination, we prepared a second, independent sequencing replicate on all PCR-negative samples with evidence for MuV or another virus, and required both replicates to contain reads matching the virus detected in the sample. We found no evidence of viruses other than MuV in PCR-positive samples.

We required the total raw read count for any genus in any sample to be twice (in practice, seven times) that in any negative control from any sequencing batch. For any sample that had one or more pathogenic viral genera that passed this filter and had de-duplicated reads well distributed across the relevant viral genome, we attempted contig assembly: we used viral-ngs to filter all sample reads against all NCBI GenBank^13^ entries matching the identified species and then *de novo* assembled reads using Trinity^31^ through viral-ngs and scaffolded against the closest matching full genome identified by a blastn query^32^. We report all viruses identified via this method in **Extended Data Table 2**.

In parallel, we used SPAdes^33^ within viral-ngs to *de novo* assemble contiguous sequence from all de-duplicated, depleted reads. We used the metagenomic tool DIAMOND v0.9.13^34^ with the nr database downloaded 29 May 2017, followed by blastn^32^ of DIAMOND-flagged contigs. Using this method, we confirmed the presence of all previously identified viruses except Influenza B virus, for which we never assembled a contiguous sequence. We found no evidence of additional viruses using this method.

### Criteria for pooling across replicates

We prepared one or more sequencing libraries from each sample and attempted to sequence and assemble a genome from each of these replicates. We required a replicate of a sample to contain 3,000 unambiguous base calls for its read data to be included in that sample’s final genome assembly. This threshold was based on the maximum number of unambiguous bases (2,820) observed in negative controls across all uncontaminated sequencing batches. One sequencing batch showed evidence of contamination: we were able to assemble 7,615 unambiguous MuV bases from a water sample, with a median coverage of 4x. For samples prepared in this batch only we implemented an additional requirement for including a replicate in pooling: the assembly must have a median coverage of at least 20x, 5 times the median coverage of the water sample.

### Multiple sequence alignment of genotype G whole genomes

We required a MuV genome to contain 11,538 unambiguous base calls (75% of the total 15,384-nucleotide genome; accession JN012242.1) for inclusion in the alignment of whole genome sequences that we used for downstream analysis. For two patients with samples taken at two time points (MuVs/Massachusetts.USA/19.16/5[G] (1) and MuVs/Massachusetts.USA/19.16/5[G] (2-20.16); MuVs/Massachusetts.USA/16.16/6[G] (1) and MuVs/Massachusetts.USA/16.16/6[G] (2-17.16)), we only included the earlier sample in downstream analyses. The final alignment of whole genome sequences contains only samples belonging to genotype G; we did not include MuVs/Massachusetts.USA/24.17/5[K], which belongs to genotype K, in the alignment.

In this alignment, we also included 25 MuV genomes published on NCBI GenBank^13^. These comprise all of the sequences with organism ‘Mumps rubulavirus’ available as of September 2017 that meet the following criteria: sequence length of at least 14,000 nucleotides, belong to genotype G, sample collection year and country of origin reported in GenBank, no evidence of extensive virus passaging or modification (for vaccine development, for example). The accession numbers are: KY969482, KY996512, KY996511, KY996510, KY680540, KY680539, KY680538, KY680537, KY006858, KY006857, KY006856, KY604739, KF738114, KF738113, KF481689, KM597072, JX287391, JX287390, JX287389, JX287387, JX287385, JN012242, JN635498, AF280799, EU370207.

We aligned MuV genomes using MAFFT v7.221^35^ with default parameters. In **Supplementary Data**, we provide the sequences and alignments used in analyses.

### Visualization of coverage depth across genomes

We plotted aggregate depth of coverage across the 200 samples whose genomes were included in the final alignment (**Extended Data Fig. 1a**) as described in ref. 36. We aligned reads against the reference genome with accession JX287389.1 and plotted over a 200-nt sliding window.

### Analysis of within- and between-sample variants

We ran V-Phaser 2.0^37^ via viral-ngs on all pooled reads mapping to a sample assembly to identify within-sample variants (**Supplementary Table 2**). To call a variant, we required a minimum of 5 forward and reverse reads, as well as no more than 10-fold strand bias, as previously described^17^. Samples with genomes generated by the sequencing batch that showed evidence of contamination (see ‘Criteria for pooling across replicates’ above) were not included in within-host variant analysis. When analyzing variants in known contacts, we used pairs of samples designated as ‘contact links’, as described in ‘Relationship between epidemiological and genetic data’ below.

Between-sample variants were called by comparing each final genome sequence to JX287385.1, the earlier of the two available whole genomes from the 2006 mumps outbreak in Iowa, USA (**Supplementary Table 2)**. We ignored all fully or partially ambiguous base calls, and excluded sequences that did not descend from the USA_2006 clade from this analysis. When examining amino acid changes in HN given vaccination status (see ‘SH and HN multiple sequence alignment’ below), we ignored sequences from patients with unknown vaccination history.

### Maximum likelihood estimation and root-to-tip regression

We generated a maximum likelihood tree using the whole genome genotype G multiple sequence alignment. We used IQ-TREE v1.3.13^38^ with a GTR substitution model and rooted the tree on the oldest sequence in this dataset (accession KF738113.1) in FigTree v1.4.2^39^.

To estimate root-to-tip distance of samples in the primary USA lineage, we subsetted the full genotype G alignment to include only samples descendent of the USA_2006 clade, including samples in this clade (see **Fig. 1a**) and used TempEst v1.5^40^ with the best fitting root according to the heuristic residual mean squared function to estimate distance from the root. We used scikit-learn^41^ in Python to perform linear regression of distances on dates.

We also generated maximum likelihood trees using the SH gene only (full 316-nucleotide mRNA), HN (coding region only), F (coding region only), and a concatenation of the aforementioned SH, HN, and F regions (**Extended Data Fig. 8**). For each tree, we started with the whole genome genotype G alignment (225 sequences) and extracted the relevant region(s). We then removed any sequence with two or more consecutive ambiguous bases (‘N’s) in any of SH, HN, or F, leaving 209 sequences in each alignment. We used IQ-TREE v1.5.5 with a GTR substitution^42^ model to generate maximum likelihood trees.

### Molecular dating using BEAST

We performed all molecular clock analyses on whole genome sequences using BEAST v1.8.4^43^. We excluded from the CDS the portion of the V protein after the insertion site^44^ because of reading frame ambiguity in that region. On the CDS, we used the SRD06 substitution model^45^, which breaks codons into two partitions (positions (1+2) and 3) with HKY substitution models^46^ and allows gamma site heterogeneity^47^ (four categories) on each. We used a separate partition on noncoding sequence with an HKY substitution model and gamma site heterogeneity. To accomodate inexact dates in seven sequences from NCBI GenBank, we used sampled tip dates^48^.

We tested six models as described in ref. 36. Each was a combination of one of two clock models (strict clock and uncorrelated relaxed clock with log-normal distribution^49^) and one of three coalescent tree priors (constant size population, exponential growth population, and Bayesian Skygrid model^50^ with 20 parameters). On each model, we estimated marginal likelihood with path sampling (PS) and stepping stone sampling (SS)^51,52^ (**Extended Data Table 3**) after sampling 100 path steps each with a chain length of 2 million.

We sampled trees and other parameters on each model by running BEAST for 200 million MCMC steps, sampling every 20,000 steps, and removing 20 million steps as burn-in. We report the mean clock rate as the substitution rate for relaxed clock models. On the sampled trees, we used TreeAnnotator v1.8.4 to find the maximum clade credibility (MCC) tree and visualized it in FigTree v1.4.3^39^. To estimate tMRCAs (**Fig. 1c, Extended Data Table 3**), we ran BEAST again for each of the six models, drawing from these same sets of sampled trees (without any parameters), for 10,000 steps, sampling every step. We selected a relaxed clock and Skygrid model for plots of tMRCA distributions (**Fig. 1c**) and MCC trees over the whole genome genotype G sequences.

### Gene- and site-specific dN/dS analyses

We used BEAST v1.8.4^43^ to estimate d*N*/d*S* per-site (**Extended Data Fig. 2a**, **Supplementary Table 3**) and per-gene (**Extended Data Fig. 2b**) using the same alignment of 225 whole genome sequences described above (again, removing the portion of the V gene after the insertion site).

For site-specific d*N*/d*S* estimation, we used the CDS as input and created a separate partition for each codon position (three partitions). We used an HKY substitution model^46^ on each partition, and an uncorrelated relaxed clock with log-normal distribution^49^ for branch rates. Here, we sampled from the same set of trees that were sampled as described above in ‘Molecular Dating using BEAST’ (relaxed clock with Skygrid tree prior). We ran BEAST for 10 million MCMC steps, sampling every 10,000 steps. We estimated site-specific d*N*/d*S* at each sampled state using renaissance counting^53,54^ and show summary statistics at each site after discarding 1 million steps as burn-in.

For per-gene estimation, we created eight separate partitions: seven correspond to the CDS of a gene (F, HN, L, M, NP, SH, partial V) and the last corresponds to noncoding sequence. For each gene partition, we used a Goldman-Yang codon model^55^ with its own parameters for d*N*/d*S* (omega) and clock rate. For the noncoding partition, we used an HKY substitution model^46^ and gamma site heterogeneity^47^ (four categories). We sampled tip dates as with the molecular clock analyses above, and used a Bayesian Skyline tree prior^56^ (10 groups). We ran BEAST for 200 million MCMC steps to sample trees and parameter values, discarded 20 million steps as burn-in, and plotted the posterior distribution of omega for each gene partition.

### Principal components analysis

The dataset for PCA consisted of all SNPs from sites with exactly two alleles in the set of all genotype G genomes. We imputed missing data with the R package missMDA^57^ and calculated principal components with the R package FactoMineR^58^. We discarded 14 samples as outliers based on visual inspection, leaving 211 samples in the final set.

### Relationship between epidemiological and genetic data

We obtained detailed epidemiological data for samples shared by MDPH from the Massachusetts Virtual Epidemiologic Network (MAVEN) surveillance system, an integrated web-based disease surveillance and case management system^59^. We defined two types of epidemiological links: ‘contact links’, between individuals who were determined to be close contacts during public health investigation and had symptom onset dates 7–33 days apart (individuals with MuV are usually considered infectious two days before through five days after onset of parotid swelling, with a typical incubation period of 16–18 days, ranging from 12–25 days)^15^; and ‘shared activity links’, between individuals who participated in the same extra-curricular activity (e.g., a sports team or university club) or frequented a specific residence or athletic facility. When we refer to epidemiological links without specifying link type, we include both types of links.

We calculated pairwise genetic distance between all pairs of samples in the whole genome genotype G alignment. For each pair, the genetic distance score is *s*/*n*, where *s* is the number of unambiguous differing sites (both sequences must have an unambiguous base at the site, and the called bases must differ) and *n* is the number of sites at which both sequences have an unambiguous base call.

To visualize the similarity between genomes and its relationship to epidemiological linkage, we performed a multidimensional scaling on sequences in clade II-outbreak (**Fig. 1a**). This clade is comprised of mostly cases from Harvard and the related community outbreak. Using their pairwise genetic distances, we calculated a metric multidimensional scaling to two dimensions in R with cmdscale^60^. We then evenly split the range of the output coordinates into a 100×100 grid and collapsed each point into this grid, and plotted the number of points at each grid coordinate; this improves visualization of nearly overlapping points (identical or near-identical genomes). We plotted curves that represent epidemiological links between cases within each of the grid coordinates. This is shown in **Extended Data Fig. 6a**.

To determine the ability of genetic distance to predict epidemiological linkage, we again looked specifically at cases within clade II-outbreak (**Fig. 1a**). Using the Python scikit-learn package^41^, we constructed a receiver operating characteristic (ROC) curve using pairwise distance between II-outbreak cases as the predictor variable and presence or absence of an epidemiological link as the binary response variable. This is shown in **Extended Data Fig. 6b**.

### Model of mumps transmission in a university setting

We developed a stochastic model for mumps transmission accounting for the natural history of infection, vaccination status, and control measures implemented in response to the outbreak at Harvard. Our stochastic model of mumps virus transmission included the stages after initial infection, the durations of which we inferred using data from previous clinical studies (**Extended Data Fig. 5a–c**). These included the gamma-distributed incubation period from infection to onset of mumps virus shedding in saliva^61^; the gamma-distributed period of latent infection from shedding onset to parotitis onset^61,62^; and the log-normally distributed time from parotitis onset to the cessation of shedding (defined in ref. 63). For asymptomatic cases, we defined the total duration of shedding (*γ*) as the sum of independent random draws from the durations of shedding before and after parotitis onset, based on the lack of any reported difference in durations of shedding for symptomatic and asymptomatic cases^61^. To account for case isolation interventions implemented at Harvard, we modeled the removal of symptomatic individuals one day after onset of parotitis. In comparison to the 70% probability for symptoms given infection among unvaccinated individuals^64^, we modeled the probability of symptoms given infection as uniformly distributed between 27.3% and 38.3%^5,16^.

We used previous estimates of the effectiveness and waning rate of mumps vaccination^5^, and of the vaccination status distribution of individuals on a university campus^65^, to account for susceptibility to infection among the Harvard population (*N*=22,000). We scaled risk for MuV infection, given exposure, to time since receipt of the last vaccine dose, yielding the hazard ratio

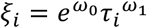

for an individual *i* who received his/her last dose *τ_i_* years previously, relative to an unvaccinated individual. For fitted values from ref. 5, estimates were below 1.0 for individuals vaccinated since 1967, when the Jeryl Lynn vaccine was introduced (**Extended Data Fig. 5d**).

Given the instantaneous hazard of infection for an as-yet uninfected individual *i* exposed to *I*(*t*) infected individuals

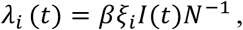

the probability of evading infection over the course of a one-day simulated time step was 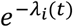. The per-contact transmission rate (*β*) was measured from the initial (pre-introduction) value of the effective reproductive number:

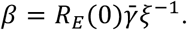

### Inferring transmission dynamics

The number of cases (71) and identification of multiple, distinct viral clades within Harvard suggested limited permeation of MuV after any introduction. We simulated dynamics of individual transmission chains to understand the epidemiological course of introduced viral lineages, and to infer values of *R_E_*(0) and the number of importations of MuV. We used the simulation model to sample from the distribution of the number of cases (*X*, including the index infection if symptomatic) resulting from a single introduction over a 1.5 year time course:

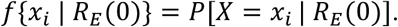

We resampled according to *f*{*x_i_*|*R_E_*(0)}to define the distribution of the cumulative number of cases (*Z*) resulting from *Y* introductions, conditioned on *R_E_*(0):

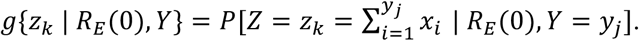

Of the 71 cases at Harvard, 66 had MuV genomes in our dataset, so we ran simulations where Z ≥ 66, drawing *k=* 66 cases at random to determine the number of distinct lineages (*S*, defined by the index infection) expected to be present within such a sample. The probability of obtaining 66 sequences, and observing *S* = *s_m_*. lineages among them, is

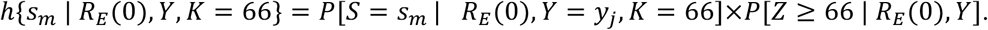

The posterior density of our model also accounted for the probability of observing 71 symptomatic cases in total. Defined in terms of the number of introductions and the initial reproductive number, the model posterior was proportional to

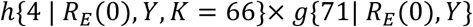

where 4 is the number of viral lineages in the 66 Harvard cases (representing clades 0-HU, II-community, and two sub-clades within clade II). We measured this probability from 100,000 iterates for each pairing of *R_E_*(0) ∊ {0.10, 0.11,…, 2.50} and *Y* ∊ {1, 2,…,200}.

Last, we defined the minimum necessary third-dose vaccine coverage (*C*) to bring the effective number below unity using the relation

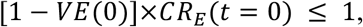

sampling from previous estimates^5^ of the effectiveness of mumps vaccination against infection, prior to waning of immunity (here defined as *VE*(0)).

### Transmission reconstruction using outbreaker

We used the R package outbreaker^66^ to reconstruct transmission for samples included in clade II-outbreak. We estimated the generation interval by fitting a gamma distribution, via maximum likelihood, to the time between symptom onset dates for cases with confirmed epidemiological links (**Extended Data Fig. 5e**, **Supplementary Data**). The resulting estimates are nearly identical to those reported in previous studies^67^ We ran outbreaker six times in parallel, each with 1 million MCMC steps, and discarded the first 10% of states as burn-in. We assessed run convergence and combined results for five of the six parallel runs to determine the reconstructed transmission tree (**Fig. 2c, right**). To reconstruct transmission using SH sequences only (**Extended Data Fig. 6c**), we extracted the SH gene from the II-outbreak alignment and ran outbreaker as described above, using the results from all six parallel runs in the analysis.

### SH and HN multiple sequence alignment

To analyze all published SH and HN MuV sequences, we searched NBCI GenBank in July 2017 for all nucleotide sequences with organism ‘Mumps rubulavirus’. We performed a pairwise alignment between each sequence *s*_i_ and a reference genome (accession: JX287389.1) using MAFFT v7.221^35^ with parameters: ‘-- localpair --maxiterate 1000 --preservecase’. We then extracted the SH sequence from each *s*_i_ based on the reference coordinates in the alignment, removing all SH sequences without the full 316-nucleotide region and all SH sequences with an insertion or deletion (‘indel’) relative to the reference. We then used MAFFT with parameters ‘--localpair --maxiterate 1000 --retree 2 --preservecase’ to create a multiple sequence alignment of the extracted SH gene sequences and removed any sequences with indels in this final alignment. We repeated the same process for the HN region, requiring the full 1749-nucleotide coding region.

In both the SH and HN alignments, we removed sequences from vaccine strains (i.e., genotype N, or another genotype marked as “(VAC)” or “vaccine”). We also removed sequences with GenBank records indicating extensive passaging. In the SH alignment only, we removed sequences with no reported collection date or country of origin, as these data are required for phylogeographic analyses. In samples with a collection decade (e.g., 1970s) but not a specific year, we assigned the first year of the decade; in samples with only a collection year, we assigned a decimal year of *year* + 0.5 (e.g., 1970.5); in samples with year and month but no day, we used the day halfway through the given month (e.g., 2015-03 becomes 2015-03-15) to calculate the decimal year; and in samples with an epidemiological week but no specific day, we approximated the decimal year as *year* + (*epi week* / 52), except samples collected in epidemiological week 52 were relabeled as week 51.999 to avoid confusion with year-only samples.

In both the SH and HN alignments, we relabeled outdated genotypes (M, E, and any sub-genotypes^68^) and constructed a maximum likelihood tree (using IQ-TREE with a GTR substitution model, as described above) to assign a genotype if one was not reported on GenBank. We preserved genotypes designated as ‘Unclassified’^68^.

To each alignment, we added all SH or HN sequences from individual patients generated in this study, except those with 2 or more consecutive ambiguous bases (‘N’s) in the SH or HN region. The sequences used in the SH and HN analyses are listed in **Supplementary Table 4**.

### SH phylogeographic analysis

To perform phylogenetic and phylogeographic analyses of the SH gene sequence, we first sampled trees using BEAST v1.8.4^43^. We used constant size population and strict clock models. Other demographic and clock models would have likely provided a better fit to the data, but it appeared that using them (especially the use of a relaxed clock) would have made it impractical to achieve convergence and sufficient sampling of trees and all parameters on the full dataset. We used the HKY substitution model^46^ with 4 rate categories and no codon partitioning. We ran BEAST in 4 replicates, each for 500 million states with sampling every 50,000 states, and removed the first 150 million states as burn-in. We verified convergence of all parameters across the 4 replicates and then combined the 4 replicates using LogCombiner.

We used TreeAnnotator to determine the maximum clade credibility (MCC) tree (**Fig. 3a**) from a resampling of 350 trees and visualized the result in FigTree v1.4.3^39^. We computed a kernel density estimate of the probability distribution of the tMRCA over all sampled states for the 11 genotypes in this dataset (**Fig. 3b**).

The full dataset has large temporal and spatial sampling biases that might affect phylogeographic results, such as estimates of migration. Using a structured coalescent to model migration and coalescent processes could alleviate these biases, but might face practical limitations on this large dataset^69^. Instead, we used resampling on the input sequences to construct distributions of estimates. To perform this resampling, we focused on only samples that were collected both within a window of time and from a geographic region with sufficient sampling. Namely, we considered only sequences sampled in 2010 or afterward and collapsed the locations shown on the full dataset (**Fig. 3a**) to just 4 regions: United States (consisting of only samples from the United States), Europe (consisting of samples whose location was labeled as Eastern Europe, Northern Europe, Southern Europe, Western Europe, or the United Kingdom), East Asia (consisting of samples whose location was labeled as Eastern Asia or Japan), and South/Southeast Asia (consisting of samples whose location was labeled as Southeast Asia or Southern Asia). We ignored samples from 5 locations: Canada; Caribbean, Central America, and South America; Middle and Eastern Africa; Northern Africa; Middle East). Then, we randomly sampled 10 sequences (without replacement) from each region for each year (i.e., 2010–2011, 2011–2012, etc.). We resampled the input sequences with this strategy 100 times.

Note that sampling biases affect this resampling strategy as well. Notably, several years in the East Asia and the South/Southeast Asia regions have fewer than 10 sequences (sometimes, zero) available for resampling (**Supplementary Table 4**), which may lead to an underestimate on the relative rates of migration involving these regions. Moreover, the high sequence similarity between SH gene sequences from United States and Europe (**Fig. 3d**) may bias upward the distributions on the relative rates of migration between these regions and on the proportion of ancestry shared between them (**Extended Data Fig. 7e**) (e.g., if there have been few true migrations between these regions, but the gene has not accumulated substitutions in the time between those migrations).

For each of the 100 resamplings of the input sequences, we ran BEAST to sample trees, as described above, for 100 million states sampling every 10,000 states; we removed 10 million states as burn-in and resampled to obtain 1,000 sampled trees.

Then, we performed phylogeographic analyses on each of the 100 samplings of input sequences by drawing from their sampled trees. We used a discrete trait substitution model^70^ on location in BEAST v1.8.4. To estimate transition rates between locations we used an nonreversible CTMC model with 4^2^ - 4 = 12 rates. Furthermore, to evaluate the significance of routes in the diffusion process, we added indicator variables to each rate through Bayesian stochastic search variable selection (BSSVS); we set the number of non-zero rates to have a Poisson prior with a mean of 3.0, placing considerable prior probability on having the fewest rates needed to explain the diffusion. We ran BEAST with 10 million states, sampling every 1,000 states, and removed the first 1 million as burn-in. At each sampled state, we logged the complete Markov jump history^71,72^, as well as a tree with the reconstructed ancestral location of each node.

To determine an MCC tree across the 100 samplings of input sequences, we ran TreeAnnotator on the sampled trees from each of the 100 samplings and then selected the MCC tree, from the 100 options, with the highest clade credibility score. We show this one, colored by reconstructed ancestral locations, in **Extended Data Fig. 7b**.

For each sampling *x*_i_ of the 100 samplings of input sequences, we counted the number of jumps between each pair of locations at each state using the complete Markov jump history after resampling the jump history every 10,000 states. For each *x*_i_, at each state, we calculated the fraction of migrations between each region pair by dividing the number of migrations between the pair by the total number of migrations at that state. To quantify support for migration routes in each *x*_i_, we calculated Bayes factors (BF) on the rate indicator variables. We calculated the posterior probability that a rate is non-zero as the mean of the indicator variable over the MCMC states, thereby providing a posterior odds. We calculated the prior probability that a rate is non-zero as the expected number of non-zero rates divided by the number of rates, which reduces to 1/*N*, where *N* is the number of locations; thus, the prior odds is 1/(*N*-1) or, in this case, 1/3. We set an upper limit of 10,000 on the BF and a lower limit of 1.0. To estimate the proportion of ancestry for each *x*_i_, we used skyline statistics via PACT^73^ to calculate proportion of ancestry at each location from each other location: in particular, for each *x*_i_, we used the sampled trees with ancestral locations as input (after resampling them every 10,000 states), padded the trees with migration events, broke the trees into temporal windows of 0.1 year going back five years prior to sampling, and estimated the proportion of history from tips in each time window.

To summarize phylogeographic results across the 100 resamplings *x*_i_ of the input sequences, we show probability distributions across the *x*_i_. When plotting the fraction of migrations to each region from each other (**Fig. 3c**), we calculated the mean of this fraction across all the sampled states in each of the 100 runs to produce a point estimate for each *x*_i_, and show the distribution of these means across the 100 *x*_i_. We calculated the Bayes factors on migration routes between the 4 regions by combining the sampled indicator variables across all 100 *x*_i_ to compute the posterior odds (**Extended Data Fig. 7c**). Similarly, the proportion of ancestry plotted between a pair of locations in a time window (**Extended Data Fig. 7d**) is the mean across the 100 *x*_i_ of the mean proportion for that pair in that time window from each *x*_i_. We calculated the pointwise percentile bands in **Extended Data Figure 7e** from the mean proportions in each *x*_i_ across the *x*_i_ (i.e., they are percentiles across the resamplings of the input sequences).

## Extended Data Figures and Tables

**Extended Data Figure 1.**
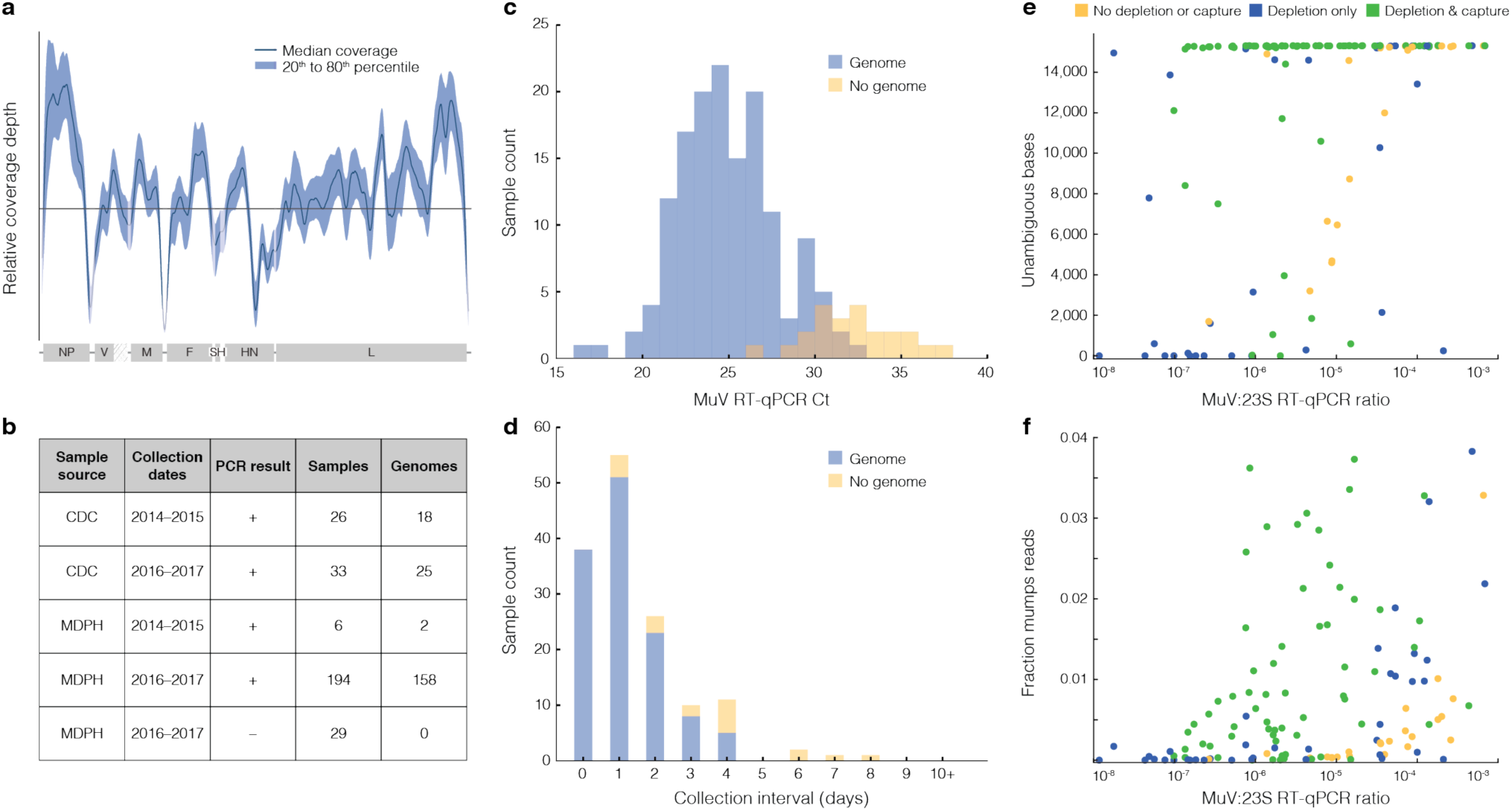
Sequencing results and predictors of outcome. (**a**) Relative sequencing depth of coverage aggregated across 200 MuV genomes. (**b**) Counts of samples received by sample source –– Massachusetts Department of Public Health (MDPH) and Centers for Disease Control and Prevention (CDC) –– and resulting genomes. (**c**) Distribution of MuV RT-qPCR cycle threshold (Ct) value, taken at sample source, for all sequencing replicates prepared with both depletion and capture (see Methods). Genome (blue): a replicate produces a genome passing the thresholds described in Methods. MuV RT-qPCR serves as a predictor of sequencing outcome. (**d**) Distribution of collection interval (days between symptom onset and sample collection) for all samples prepared with both depletion and capture. Genome (blue) is defined as in (c). Samples taken more than 4 days after symptom onset did not produce genomes in this study^74^. (**e**) Number of unambiguous bases in the genome assembly of each sample by MuV:23S ratio (MuV copies by MuV RT-qPCR divided by 23S copies by 23S RT-qPCR; see Methods). Each point is a replicate, colored by sequencing preparation method. (**f**) Normalized MuV reads (unique MuV reads divided by raw sequencing depth) in each sample by MuV:23S ratio. Points are as in (d). Nine points with fraction mumps reads >0.04 are not shown. In (c–f), reads from each replicate were downsampled to 1 million prior to assembly (see Methods). In (e) and (f), 1 point with a MuV:23S ratio <10^−8^ and 3 points with a MuV:23S ratio >10^−3^ are not shown.

**Extended Data Figure 2.**
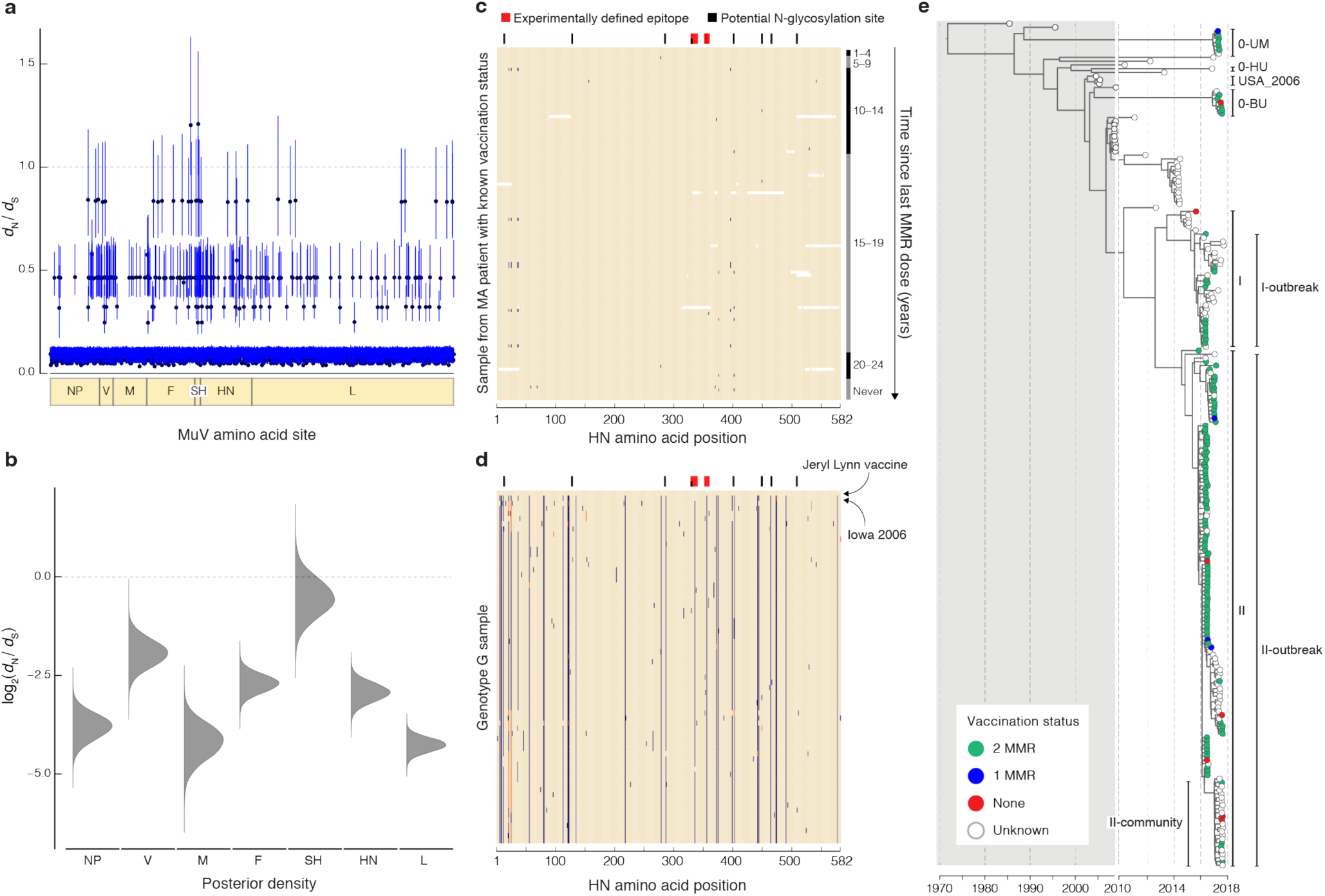
Amino acid substitution in the MuV genome. (**a**) Estimate of dN/dS at each amino acid site in MuV coding regions, calculated across all 225 genotype G genomes used in this study. At each site, the mean estimate and 95% credible interval is shown. (**b**) Posterior density of dN/dS in each MuV gene, using the same dataset. (**c**) Variation in genomes generated in this study. Each row represents one of the 119 MuV HN amino acid sequences from the individuals in our study who had known vaccine status. Samples are displayed in order of descending time since last MMR vaccine dose. Colored variants indicate variation from the consensus of all included sequences. (**d**) Variation in all published genotype G HN sequences. Each row represents one of the 456 publicly available MuV genotype G HN sequences (including from genomes generated in this study). Identical sequences are collapsed and then grouped by hierarchical clustering. In both panels, amino acid substitutions relative to the Jeryl Lynn vaccine strain are highlighted in blue, with orange indicating a second variant allele and green indicating a third. Red bars indicate experimentally-identified neutralizing antibody epitopes, and black bars indicate potential N-glycosylation sites. (**e**) Maximum clade credibility tree of the 225 MuV genotype G genomes used in this study, colored by vaccination status. Clades are labeled as in **Fig. 1a**.

**Extended Data Figure 3.**
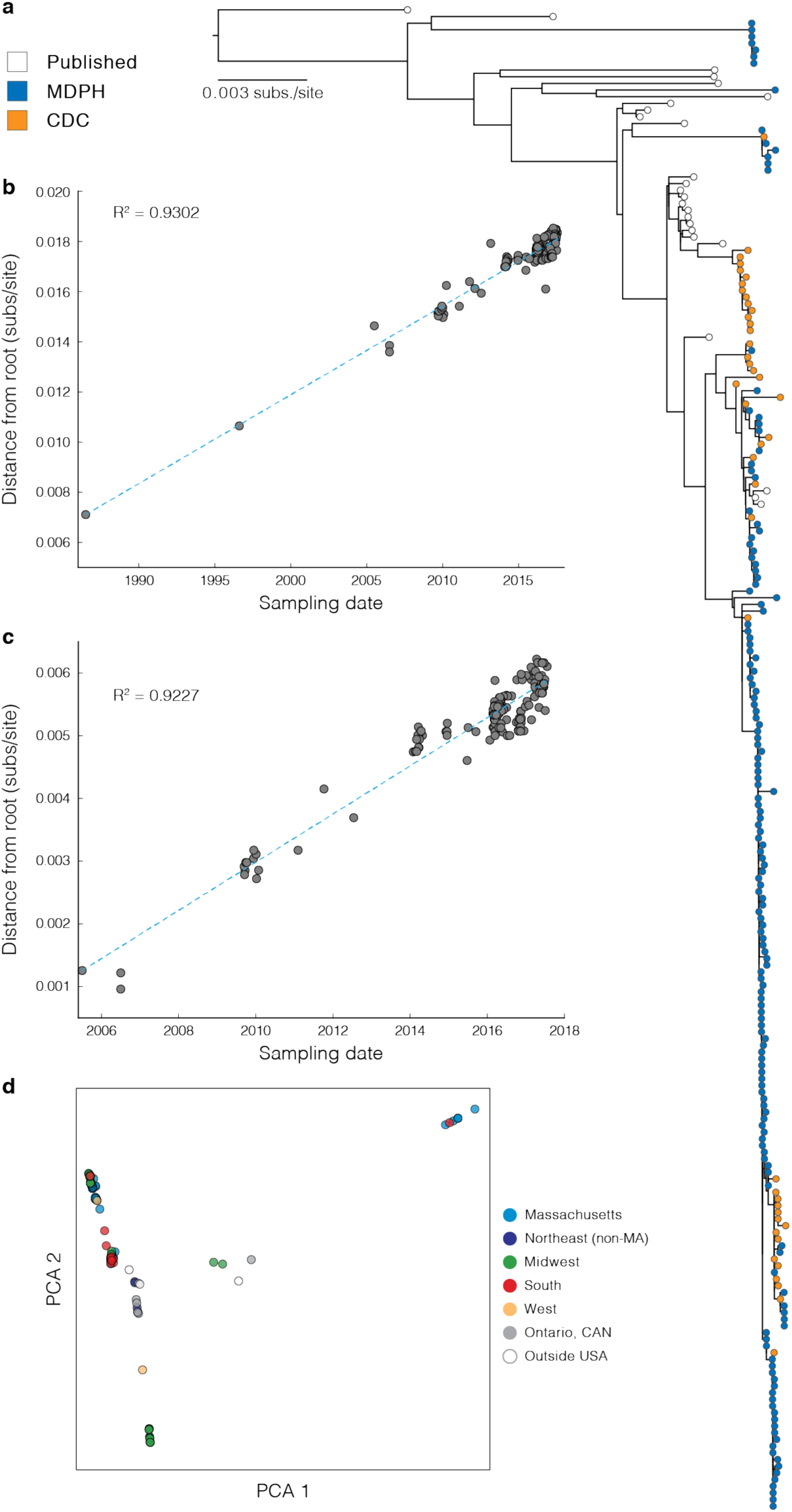
Maximum likelihood tree, root-to-tip regression, and principal components analysis. (**a**) Maximum likelihood tree of the 225 MuV genotype G genomes used in this study. Tips are colored by sample source (MDPH or CDC); previously-published genomes are indicated by unfilled circles. (**b**) Root-to-tip regression of genomes shown in (a), rooted on GenBank accession KF738113 (Pune.IND, 1986). (**c**) Root-to-tip regression of genomes in the clade containing the two USA 2006 sequences (USA_2006; see **Fig. 1**), as well as their descendents. (**d**) Principal components analysis of genetic variants from the genomes in (a). Each point is a genome colored by its geographic location.

**Extended Data Figure 4.**
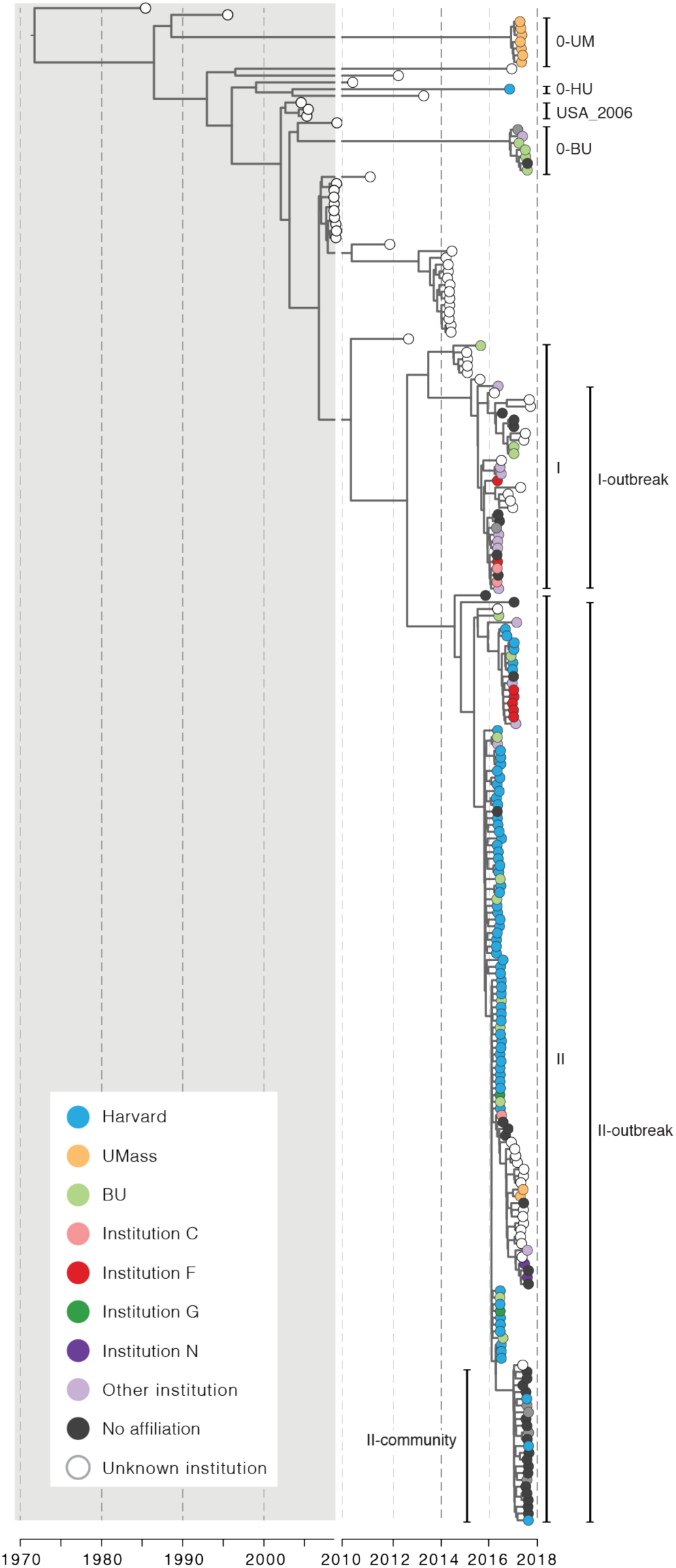
Phylogenetic tree colored by institution. Maximum clade credibility tree of the 225 MuV genotype G genomes used in this study, colored by academic institution. Clades are labeled as in **Fig. 1a**.

**Extended Data Figure 5.**
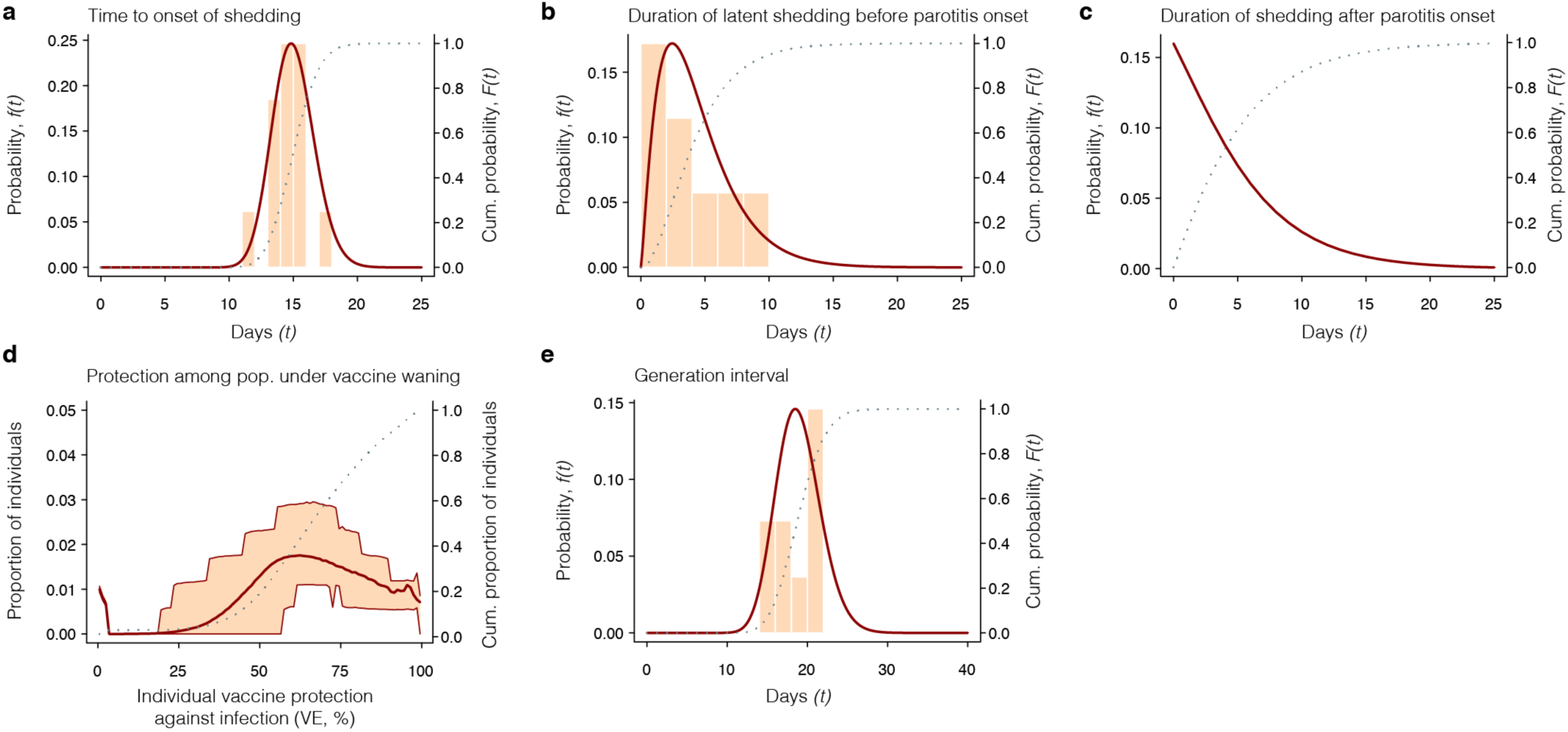
Parameters used in epidemiological models. We illustrate fitted distributions of parameters of the modeled natural history of mumps infection. (**a**) We calibrate a gamma distribution to the duration of the incubation period—defined from the time of mumps virus exposure to the onset of shedding—using data from experimental human mumps virus infections with known exposure times^61^. (**b**) Onset of mumps shedding generally precedes onset of symptoms in the clinical course. We fit a gamma distribution describing the period of latent shedding to pooled data from two studies^61,62^, and (**c**) apply previous estimates of the distribution of the duration of shedding after parotitis onset^63^. (**d**) We obtain estimates of the distribution of vaccine protection within a university protection by pairing previous estimates of the association between the strength of vaccine protection and time since receipt of the last dose^5^ to data on vaccine coverage in a large university^65^. (**e**) We infer the distribution of the generation interval length in the Harvard using data from 10 cases with known exposure sources (‘contact link’). A gamma distribution fitted by maximum likelihood recovers mean and dispersion estimates nearly identical to those reported in earlier mumps outbreaks^67^.

**Extended Data Figure 6.**
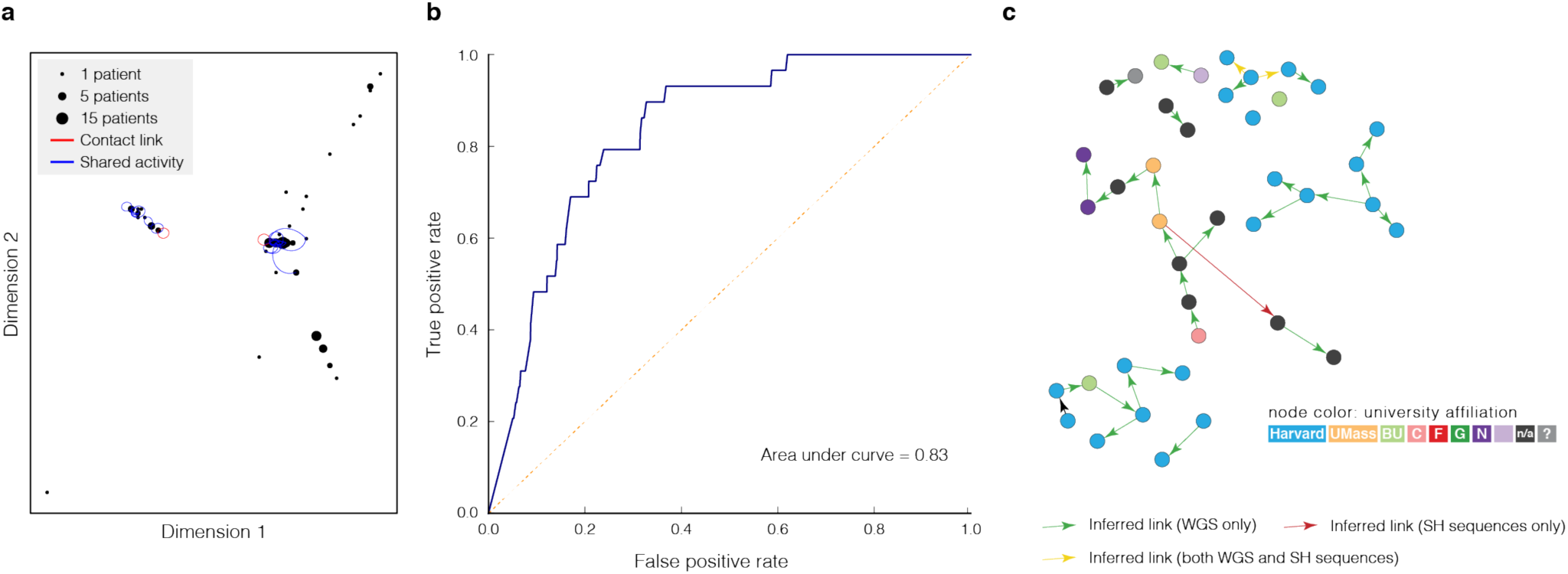
Connection between epidemiological and genetic data. (**a**) Multidimensional scaling applied to samples in clade II-outbreak (see **Fig. 1a**). Each point is a MuV genome and pairwise dissimilarities are based on Hamming distance (see Methods). Genomes with known epidemiological links are connected with a red line. (**b**) Receiver operating characteristic (ROC) curve for samples within clade II-outbreak using pairwise genetic distance (calculated as in (a)) as a predictor of epidemiological linkage. (**c**) Transmission reconstruction using individuals within clade II-outbreak, using the SH gene or whole genome sequences. Inferred links with probability **≥**0.7 are shown; arrows are colored by dataset. Nodes are colored by institution, and nodes with no inferred links are not shown. See also **Fig. 2c, right**.

**Extended Data Figure 7.**
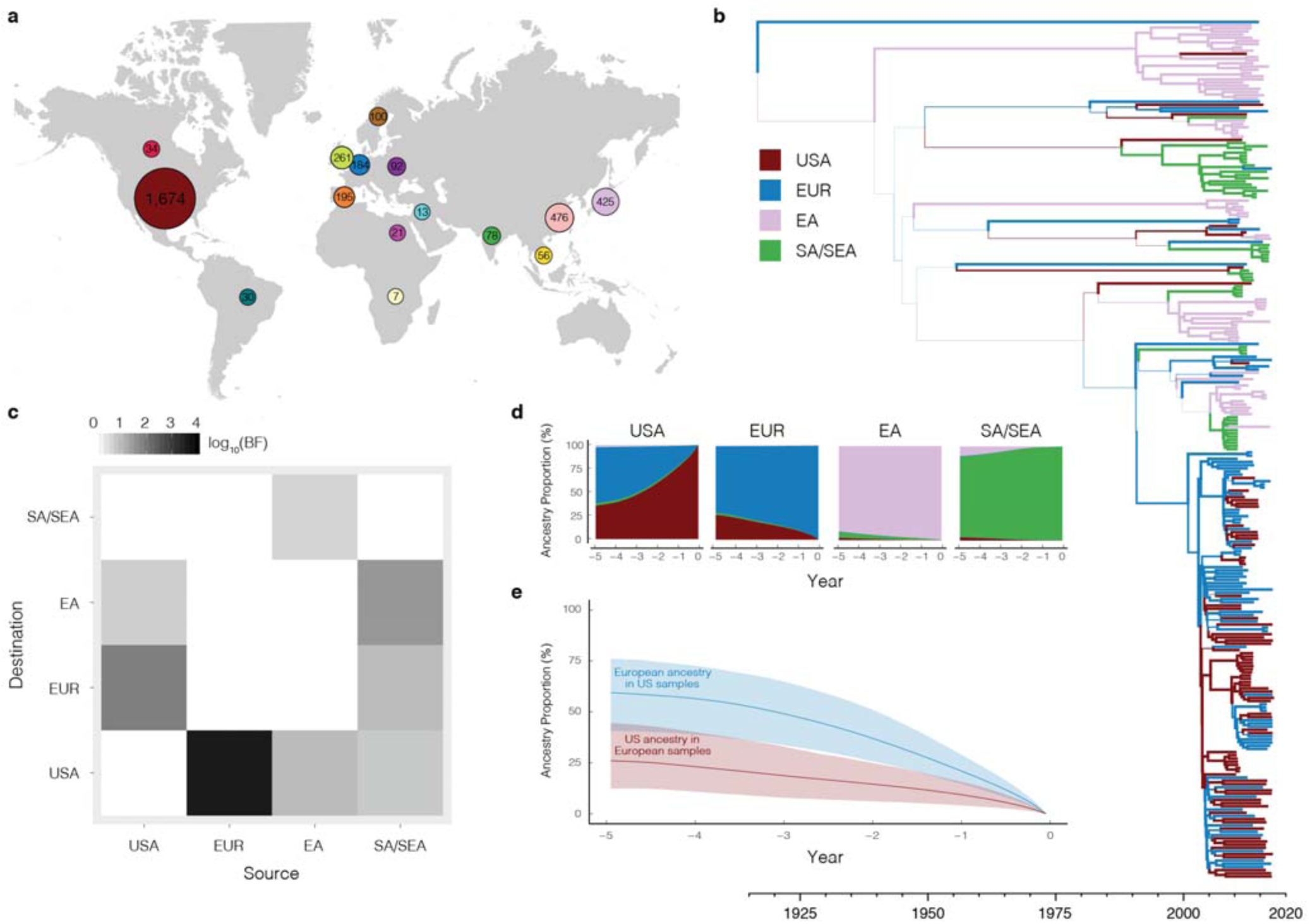
Additional analyses of global MuV spread using SH gene sequences. (**a**) World map indicating number of SH sequences in our dataset from each of 15 regions. (**b**) Tree with the highest clade credibility across all trees generated on resampled input from 4 world regions: South Asia/Southeast Asia (SA/SEA), East Asia (EA), Europe (EUR), and United States (USA) (see Methods for details regarding geographic and temporal resampling of sequences). Branch line thickness corresponds to posterior support for ancestry (indicated by branch color). (**c**) Migration between the 4 regions shown in (a). Shading of each migration route indicates its statistical support (quantified with Bayes factors) in explaining the diffusion of MuV. (**d**) Average proportion of geographic ancestry of samples in each of the 4 world regions (labeled) from each of the 4 regions (colored), going back 5 years from sample collection. Colors are as in (b). (**e**) Average proportion of EUR in geographic ancestry of USA samples, and vice-versa. Shaded regions are pointwise percentile bands (2.5% to 97.5%) across 100 resamplings of the input sequences.

**Extended Data Figure 8.**
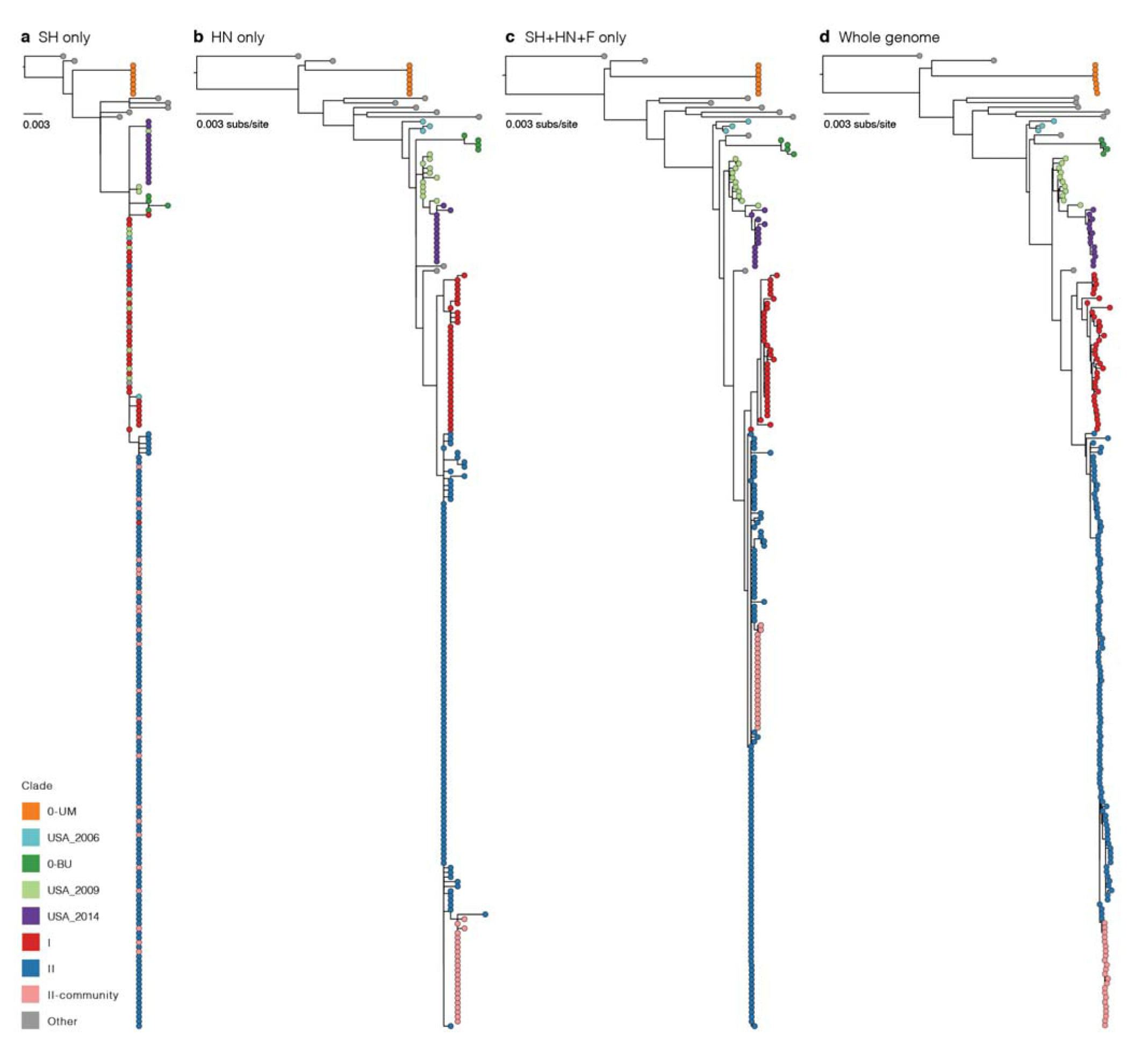
Trees produced with single- and multi-gene sequences. Maximum likelihood trees using (**a**) the SH gene only, (**b**) the HN protein only, (**c**) a concatenation of the HN protein, the fusion protein (F) coding region, and the SH gene, and (**d**) the complete MuV genome. In all panels, tips are colored by clades as defined in **Fig. 1a** and **Extended Data Table 3**. The HN protein sequence does a significantly better job at capturing the epidemiologically-relevant clades than the SH gene, and the tree created from SH+HN+F (nearly 25% of the genome) closely resembles the tree created from whole genome sequences.

**Extended Data Table 1.** Sample metadata. (**a**) Demographic information of all MuV cases in MA between 2014-01-01 and 2017-06-30, and the subset of these included in this study. (**b**) Relevant metadata for all samples PCR-negative for MuV collected during the same time period, 29 of which we attempted to sequence.

|  |  | All cases (n=375) | Study cases (n=198) |
| --- | --- | --- | --- |
| Gender | Male | 49.33% | 49.50% |
|  | Female | 50.67% | 50.50% |
| Age | <5 years | 1.87% | 0.00% |
|  | 5–9 years | 1.07% | 0.00% |
|  | 10–14 years | 0.27% | 0.51% |
|  | 15–19 years | 21.87% | 23.23% |
|  | 20–24 years | 42.40% | 48.48% |
|  | 25–29 years | 12.27% | 11.62% |
|  | 30+ years | 20.27% | 16.16% |
| Vaccination status | 2+ MMR | 64.27% | 64.65% |
|  | 1 MMR | 4.53% | 2.53% |
|  | Unvaccinated | 4.53% | 5.56% |
|  | Unknown | 26.67% | 27.27% |

**Extended Data Table 1. Sample metadata.** (a) Demographic information of all MuV cases in MA between 2014-01-01 and 2017-06-30, and the subset of these included in this study.
|  |  | Negatives (n=521) |
| --- | --- | --- |
| Vaccination status | 2+ MMR | 47.60% |
|  | 1 MMR | 7.87% |
|  | Unvaccinated | 3.84% |
|  | Unknown | 40.69% |
| Time since vaccination | <1 year | 2.69% |
|  | 1–4 years | 9.21% |
|  | 5–9 years | 7.10% |
|  | 10–14 years | 11.71% |
|  | 15–19 years | 18.43% |
|  | 20–24 years | 4.41% |
|  | 25+ years | 1.92% |
|  | Unvaccinated | 3.84% |
|  | Unknown | 40.69% |
| Time to collection | 0 days | 18.04% |
|  | 1 day | 30.71% |
|  | 2 days | 19.96% |
|  | 3 days | 9.79% |
|  | 4 days | 6.33% |
|  | 5+ days | 6.72% |
|  | Unknown | 8.45% |

**Extended Data Table 2.** Viruses identified in mumps PCR-negative samples. Influenzavirus B is the only multipartite virus listed, and we identify 51 reads mapping to 6 of the 8 segments: in order, 2, 6, 0, 0, 8, 10, 4, 21 reads to each segment.

| Pathogen | Sample | Raw reads | Taxon-filtered reads | Contigs | Unambiguous bases | Median coverage | Genome coverage |
| --- | --- | --- | --- | --- | --- | --- | --- |
| Human Parainfluenza Virus 3 | MA/24.16/NEG-1 | 14,455,928 | 2,560 | 6 | 13,859 | 8 | 89.6% |
| Human Parainfluenza Virus 2 | MA/39.16/NEG-1 | 12,327,238 | 294 | 5 | 210 | 0 | 1.3% |
| Human Coronavirus OC43 | MA/50.16/NEG-1 | 35,708,954 | 46,136 | 12 | 1,976 | 1 | 6.5% |
| Influenzavirus B | MA/10.16/NEG-1 | 15,280,400 | 54* | 0 | — | — | — |
| Mumps rubulavirus | MA/9.16/NEG-2 | 4,293,704 | 344 | 6 | 307 | 1 | 2.0% |

**Extended Data Table 3.**
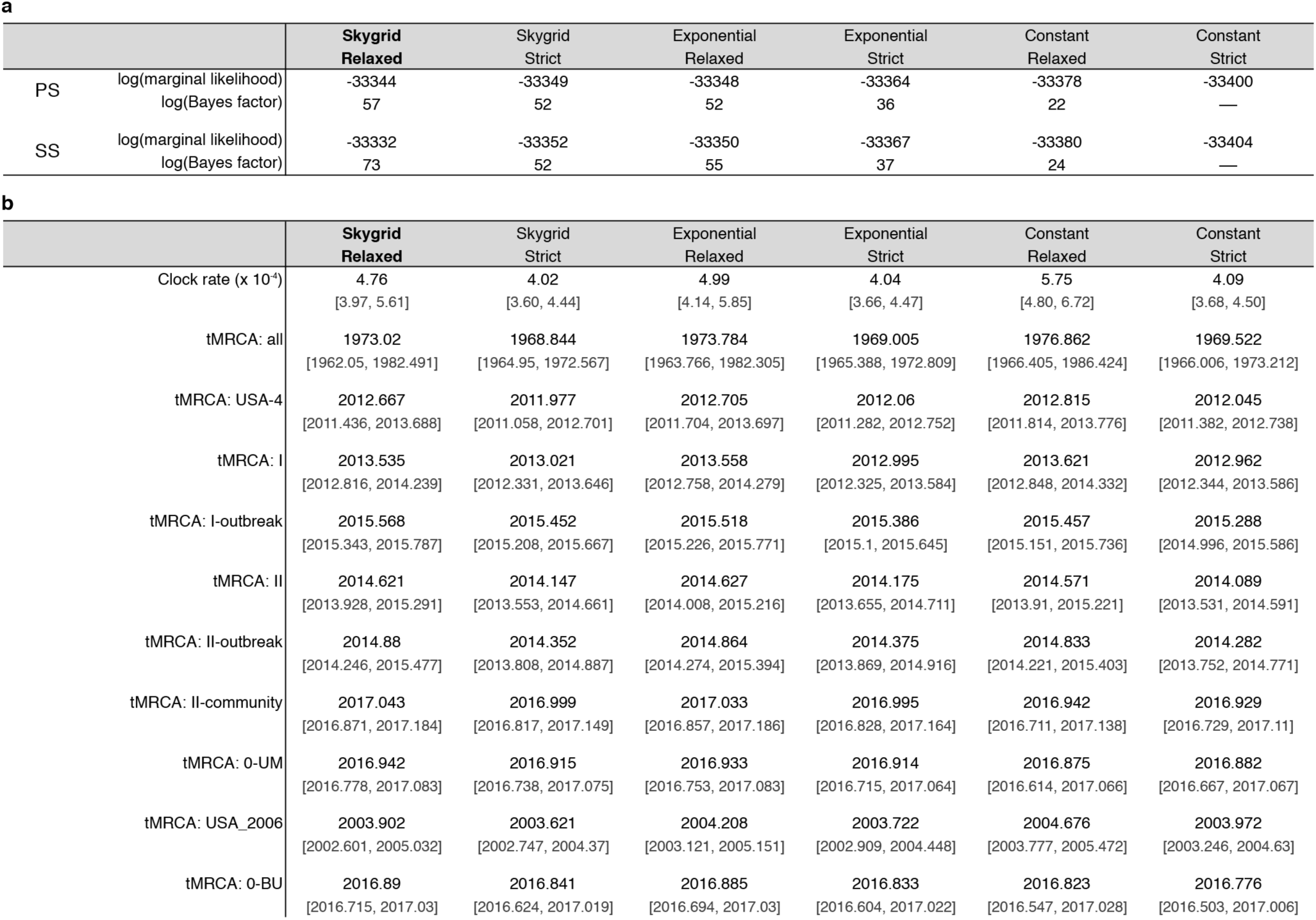
Model selection and tMRCA estimates across models. (**a**) Marginal likelihoods estimated in 6 models: combinations of three coalescent tree priors (constant size population, exponential growth population, and Skygrid) and two clock models (strict clock and uncorrelated relaxed clock with log-normal distribution). Estimates are with path-sampling (PS) and stepping-stone sampling (SS). The Bayes factors are calculated against the model with constant size population and a strict clock. (**b**) Mean estimates of clock rate, date of tree root, and tMRCAs of the clades shown in **Fig. 1a** (excluding clade 0- HU, which consists of one sample). USA-4 corresponds to ‘Clades I and II’ in **Fig. 1**. Below each mean estimate is the 95% highest posterior density interval. We calculated tMRCAs of additional unlabeled nodes (see **Supplementary Data**).

**a**
|  |  | Skygrid Relaxed | Skygrid Strict | Exponential Relaxed | Exponential Strict | Constant Relaxed | Constant Strict |
| --- | --- | --- | --- | --- | --- | --- | --- |
| PS | log(marginal likelihood) | -33344 | -33349 | -33348 | -33364 | -33378 | -33400 |
|  | log(Bayes factor) | 57 | 52 | 52 | 36 | 22 | — |
| SS | log(marginal likelihood) | -33332 | -33352 | -33350 | -33367 | -33380 | -33404 |
|  | log(Bayes factor) | 73 | 52 | 55 | 37 | 24 | — |

**Extended Data Table 3. Model selection and tMRCA estimates across models.**
|  |  | Skygrid Relaxed | Skygrid Strict | Exponential Relaxed | Exponential Strict | Constant Relaxed | Constant Strict |
| --- | --- | --- | --- | --- | --- | --- | --- |
| Clock rate (x 10<sup>-4</sup>) |  | 4.76 | 4.02 | 4.99 | 4.04 | 5.75 | 4.09 |
|  |  | [3.97, 5.61] | [3.60, 4.44] | [4.14, 5.85] | [3.66, 4.47] | [4.80, 6.72] | [3.68, 4.50] |
| tMRCA: all |  | 1973.02 | 1968.844 | 1973.784 | 1969.005 | 1976.862 | 1969.522 |
|  |  | [1962.05, 1982.491] | [1964.95, 1972.567] | [1963.766, 1982.305] | [1965.388, 1972.809] | [1966.405, 1986.424] | [1966.006, 1973.212] |
| tMRCA: USA-4 |  | 2012.667 | 2011.977 | 2012.705 | 2012.06 | 2012.815 | 2012.045 |
|  |  | [2011.436, 2013.688] | [2011.058, 2012.701] | [2011.704, 2013.697] | [2011.282, 2012.752] | [2011.814, 2013.776] | [2011.382, 2012.738] |
| tMRCA: I |  | 2013.535 | 2013.021 | 2013.558 | 2012.995 | 2013.621 | 2012.962 |
|  |  | [2012.816, 2014.239] | [2012.331, 2013.646] | [2012.758, 2014.279] | [2012.325, 2013.584] | [2012.848, 2014.332] | [2012.344, 2013.586] |
| tMRCA: I-outbreak |  | 2015.568 | 2015.452 | 2015.518 | 2015.386 | 2015.457 | 2015.288 |
|  |  | [2015.343, 2015.787] | [2015.208, 2015.667] | [2015.226, 2015.771] | [2015.1, 2015.645] | [2015.151, 2015.736] | [2014.996, 2015.586] |
| tMRCA: II |  | 2014.621 | 2014.147 | 2014.627 | 2014.175 | 2014.571 | 2014.089 |
|  |  | [2013.928, 2015.291] | [2013.553, 2014.661] | [2014.008, 2015.216] | [2013.655, 2014.711] | [2013.91, 2015.221] | [2013.531, 2014.591] |
| tMRCA: II-outbreak |  | 2014.88 | 2014.352 | 2014.864 | 2014.375 | 2014.833 | 2014.282 |
|  |  | [2014.246, 2015.477] | [2013.808, 2014.887] | [2014.274, 2015.394] | [2013.869, 2014.916] | [2014.221, 2015.403] | [2013.752, 2014.771] |
| tMRCA: II-community |  | 2017.043 | 2016.999 | 2017.033 | 2016.995 | 2016.942 | 2016.929 |
|  |  | [2016.871, 2017.184] | [2016.817, 2017.149] | [2016.857, 2017.186] | [2016.828, 2017.164] | [2016.711, 2017.138] | [2016.729, 2017.11] |
| tMRCA: 0-UM |  | 2016.942 | 2016.915 | 2016.933 | 2016.914 | 2016.875 | 2016.882 |
|  |  | [2016.778, 2017.083] | [2016.738, 2017.075] | [2016.753, 2017.083] | [2016.715, 2017.064] | [2016.614, 2017.066] | [2016.667, 2017.067] |
| tMRCA: USA_2006 |  | 2003.902 | 2003.621 | 2004.208 | 2003.722 | 2004.676 | 2003.972 |
|  |  | [2002.601, 2005.032] | [2002.747, 2004.37] | [2003.121, 2005.151] | [2002.909, 2004.448] | [2003.777, 2005.472] | [2003.246, 2004.63] |
| tMRCA: 0-BU |  | 2016.89 | 2016.841 | 2016.885 | 2016.833 | 2016.823 | 2016.776 |
|  |  | [2016.715, 2017.03] | [2016.624, 2017.019] | [2016.694, 2017.03] | [2016.604, 2017.022] | [2016.547, 2017.028] | [2016.503, 2017.006] |

## List of Supplementary Tables

**Supplementary Table 1.** Information and metadata regarding all mumps PCR-positive and PCR-negative samples on which sequencing was attempted.

**Supplementary Table 2.** List of identified nonsynonymous single nucleotide polymorphisms (SNPs), including the frequency of each SNP. Separate list of identified within-host variants above 2% frequency.

**Supplementary Table 3.** Calculated d*N*/d*S* value and confidence interval for each amino acid position across the MuV genome.

**Supplementary Table 4.** List of SH and HN sequences and corresponding metadata used in analysis, as well as SH sequences available for resampling.

## References

1. Centers for Disease Control and Prevention. Mumps Cases and Outbreaks. (2018). Available at: https://www.cdc.gov/mumps/outbreaks.html. (Accessed: 9th March 2018)

2. Centers for Disease Control and Prevention. FastStats - Immunization. J. Infect. Dis. 198, 508–515 (2017).

3. Centers for Disease Control and Prevention. Measles Prevention: Recommendations of the Immunization Practices Advisory Committee (ACIP). MMWR Surveill. Summ. 38,1–18 (1989).

4. Dayan, G. H. et al. Recent resurgence of mumps in the United States. N. Engl. J. Med. 358, 1580–1589 (2008).

5. Lewnard, J. A. & Grad, Y. H. Vaccine waning and mumps re-emergence in the United States. Sci. Transl. Med. (2018).

6. Centers for Disease Control and Prevention. National Notifiable Diseases Surveillance System. (2017).

7. Barskey, A. E. et al. Mumps outbreak in Orthodox Jewish communities in the United States. N. Engl. J. Med. 367, 1704–1713 (2012).

8. Matranga, C. B. et al. Enhanced methods for unbiased deep sequencing of Lassa and Ebola RNA viruses from clinical and biological samples. Genome Biol. 15, (2014).

9. Siddle, K. J. et al. Capturing diverse microbial sequence with comprehensive and scalable probe design. BioRxiv (2018). doi:10.1101/279570

10. Barrabeig, I. et al. Viral etiology of mumps-like illnesses in suspected mumps cases reported in Catalonia, Spain. Hum. Vaccin. Immunother. 11, 282–287 (2015).

11. Davidkin, I., Jokinen, S., Paananen, A., Leinikki, P. & Peltola, H. Etiology of mumps-like illnesses in children and adolescents vaccinated for measles, mumps, and rubella. J. Infect. Dis. 191, 719–723 (2005).

12. Centers for Disease Control and Prevention. Influenza & Parotitis: Question & Answers for Health Care Providers. Cell 161, 1516–1526 (2016).

13. Clark, K., Karsch-Mizrachi, I., Lipman, D. J., Ostell, J. & Sayers, E. W. GenBank. Nucleic Acids Res. 44, D67 (2016).

14. Wolinsky, J. S., Waxham, M. N. & Server, A. C. Protective effects of glycoprotein-specific monoclonal antibodies on the course of experimental mumps virus meningoencephalitis. J. Virol. 53, 727–734 (1985).

15. Kimberlin, D. W., Brady, M. T., Jackson, M. A. & Long, S. S. Red Book, (2015): 2015 Report of the Committee on Infectious Diseases. (Am Acad Pediatrics, 2015).

16. Dittrich, S. et al. Assessment of serological evidence for mumps virus infection in vaccinated children. Vaccine 29, 9271–9275 (2011).

17. Gire, S. K. et al. Genomic surveillance elucidates Ebola virus origin and transmission during the 2014 outbreak. Science 345, 1369–1372 (2014).

18. Park, D. J. et al. Ebola Virus Epidemiology, Transmission, and Evolution during Seven Months in Sierra Leone. Cell 161, 1516–1526 (2015).

19. Didelot, X., Gardy, J. & Colijn, C. Bayesian inference of infectious disease transmission from whole-genome sequence data. Mol. Biol. Evol. 31, 1869–1879 (2014).

20. Yeo, R. P., Afzal, M. A., Forsey, T. & Rima, B. K. Identification of a new mumps virus lineage by nucleotide sequence analysis of the SH gene of ten different strains. Arch. Virol. 128, 371–377 (1993).

21. Jin, L. et al. Proposal for genetic characterisation of wild-type mumps strains: preliminary standardisation of the nomenclature. Arch. Virol. 150, 1903–1909 (2005).

22. Jin, L. et al. Genomic diversity of mumps virus and global distribution of the 12 genotypes. Rev. Med. Virol. 25, 85–101 (2015).

23. Cui, A. et al. Evolutionary analysis of mumps viruses of genotype F collected in mainland China in 2001-2015. Sci. Rep. 7, 17144 (2017).

24. Marin, M., Marlow, M., Moore, K. L. & Patel, M. Recommendation of the Advisory Committee on Immunization Practices for Use of a Third Dose of Mumps Virus--Containing Vaccine in Persons at Increased Risk for Mumps During an Outbreak. MMWR Morb. Mortal. Wkly. Rep. 67, 33–38 (2018).

25. Centers for Disease Control and Prevention. Mumps 2012 Case Definition. National Notifiable Diseases Surveillance System (NNDSS) (2012). Available at: https://wwwn.cdc.gov/nndss/conditions/mumps/case-definition/2012/.

26. Centers for Disease Control and Prevention. Real-time (TaqMan) RT-PCR Assay for the Detection of Mumps Virus RNA in Clinical Samples. (2010). Available at: https://www.cdc.gov/mumps/downloads/lab-rt-pcr-assay-detect.pdf.

27. Van Camp, G., Chapelle, S. & De Wachter, R. Amplification and sequencing of variable regions in bacterial 23S ribosomal RNA genes with conserved primer sequences. Curr. Microbiol. 27, 147–151 (1993).

28. Chris Tomkins-Tinch, Simon Ye, Hayden Metsky, Irwin Jungreis, Rachel Sealfon, Xiao Yang, Kristian Andersen, Mike Lin, and Daniel Park. *viral-ngs*.

29. Li, H. et al. The Sequence Alignment/Map format and SAMtools. Bioinformatics 25, 2078–2079 (2009).

30. Wood, D. E. & Salzberg, S. L. Kraken: ultrafast metagenomic sequence classification using exact alignments. Genome Biol. 15, R46 (2014).

31. Grabherr, M. G. et al. Full-length transcriptome assembly from RNA-Seq data without a reference genome. Nat. Biotechnol. 29, 644–652 (2011).

32. Altschul, S. F., Gish, W., Miller, W., Myers, E. W. & Lipman, D. J. Basic local alignment search tool. J. Mol. Biol. 215, 403–410 (1990).

33. Bankevich, A. et al. SPAdes: a new genome assembly algorithm and its applications to single-cell sequencing. J. Comput. Biol. 19, 455–477 (2012).

34. Buchfink, B., Xie, C. & Huson, D. H. Fast and sensitive protein alignment using DIAMOND. Nat. Methods 12, 59–60 (2015).

35. Katoh, K. & Standley, D. M. MAFFT multiple sequence alignment software version 7: improvements in performance and usability. Mol. Biol. Evol. 30, 772–780 (2013).

36. Metsky, H. C. et al. Zika virus evolution and spread in the Americas. Nature 546, 411–415 (2017).

37. Yang, X., Charlebois, P., Macalalad, A., Henn, M. R. & Zody, M. C. V-Phaser 2: variant inference for viral populations. BMC Genomics 14, 674 (2013).

38. Nguyen, L.-T., Schmidt, H. A., von Haeseler, A. & Minh, B. Q. IQ-TREE: a fast and effective stochastic algorithm for estimating maximum-likelihood phylogenies. Mol. Biol. Evol. 32, 268–274 (2015).

39. Rambaut, A. FigTree. Edinburgh, UK: Inst. Evol. Biol., Univ. Edinburgh. (2016). Available at: http://tree.bio.ed.ac.uk/software/figtree/.

40. Rambaut, A., Lam, T. T., Max Carvalho, L. & Pybus, O. G. Exploring the temporal structure of heterochronous sequences using TempEst (formerly Path-O-Gen). Virus Evol 2, vew007 (2016).

41. Pedregosa, F. et al. Scikit-learn: Machine Learning in Python. J. Mach. Learn. Res. 12, 2825–2830 (2011).

42. Tavaré, S. Some probabilistic and statistical problems in the analysis of DNA sequences. Lectures on mathematics in the life sciences 17, 57–86 (1986).

43. Drummond, A. J., Suchard, M. A., Xie, D. & Rambaut, A. Bayesian phylogenetics with BEAUti and the BEAST 1.7. Mol. Biol. Evol. 29, 1969–1973 (2012).

44. Paterson, R. G. & Lamb, R. A. RNA editing by G-nucleotide insertion in mumps virus P-gene mRNA transcripts. J. Virol. 64, 4137–4145 (1990).

45. Shapiro, B., Rambaut, A. & Drummond, A. J. Choosing appropriate substitution models for the phylogenetic analysis of protein-coding sequences. Mol. Biol. Evol. 23, 7–9 (2006).

46. Hasegawa, M., Kishino, H. & Yano, T. Dating of the human-ape splitting by a molecular clock of mitochondrial DNA. J. Mol. Evol. 22, 160–174 (1985).

47. Yang, Z. Maximum likelihood phylogenetic estimation from DNA sequences with variable rates over sites: approximate methods. J. Mol. Evol. 39, 306–314 (1994).

48. Shapiro, B. et al. A Bayesian phylogenetic method to estimate unknown sequence ages. Mol. Biol. Evol. 28, 879–887 (2011).

49. Drummond, A. J., Ho, S. Y. W., Phillips, M. J. & Rambaut, A. Relaxed phylogenetics and dating with confidence. PLoS Biol. 4, e88 (2006).

50. Gill, M. S. et al. Improving Bayesian population dynamics inference: a coalescent-based model for multiple loci. Mol. Biol. Evol. 30, 713–724 (2013).

51. Baele, G. et al. Improving the accuracy of demographic and molecular clock model comparison while accommodating phylogenetic uncertainty. Mol. Biol. Evol. 29, 2157–2167 (2012).

52. Baele, G., Li, W. L. S., Drummond, A. J., Suchard, M. A. & Lemey, P. Accurate model selection of relaxed molecular clocks in bayesian phylogenetics. Mol. Biol. Evol. 30, 239–243 (2013).

53. O’Brien, J. D., Minin, V. N. & Suchard, M. A. Learning to count: robust estimates for labeled distances between molecular sequences. Mol. Biol. Evol. 26, 801–814 (2009).

54. Lemey, P., Minin, V. N., Bielejec, F., Kosakovsky Pond, S. L. & Suchard, M. A. A counting renaissance: combining stochastic mapping and empirical Bayes to quickly detect amino acid sites under positive selection. Bioinformatics 28, 3248–3256 (2012).

55. Goldman, N. & Yang, Z. A codon-based model of nucleotide substitution for protein-coding DNA sequences. Mol. Biol. Evol. 11, 725–736 (1994).

56. Drummond, A. J., Rambaut, A., Shapiro, B. & Pybus, O. G. Bayesian coalescent inference of past population dynamics from molecular sequences. Mol. Biol. Evol. 22, 1185–1192 (2005).

57. Josse, J., Husson, F. & Others. missMDA: a package for handling missing values in multivariate data analysis. J. Stat. Softw. 70, 1–31 (2016).

58. Lê, S., Josse, J. & Husson, F. FactoMineR: an R package for multivariate analysis. J. Stat. Softw. (2008).

59. Troppy, S., Haney, G., Cocoros, N., Cranston, K. & DeMaria, A., Jr. Infectious disease surveillance in the 21st century: an integrated web-based surveillance and case management system. Public Health Rep. 129, 132–138 (2014).

60. R Core Team. R: A Language and Environment for Statistical Computing. R Foundation for Statistical Computing (2016). Available at: https://www.R-project.org/.

61. Henle, G. & Henle, W. Isolation of mumps virus from human beings with induced apparent or inapparent infections. J. Exp. Med. 88, 223–232 (1948).

62. Ennis, F. A. & Jackson, D. Isolation of virus during the incubation period of mumps infection. J. Pediatr. 72, 536–537 (1968).

63. Polgreen, P. M. et al. The duration of mumps virus shedding after the onset of symptoms. Clin. Infect. Dis. 46, 1447–1449 (2008).

64. Galazka, A. M., Robertson, S. E. & Kraigher, A. Mumps and mumps vaccine: a global review. Bull. World Health Organ. 77, 3–14 (1999).

65. Cardemil, C. V. et al. Effectiveness of a Third Dose of MMR Vaccine for Mumps Outbreak Control. N. Engl. J. Med. 377, 947–956 (2017).

66. Jombart, T. et al. Bayesian reconstruction of disease outbreaks by combining epidemiologic and genomic data. PLoS Comput. Biol. 10, e1003457 (2014).

67. Vink, M. A., Bootsma, M. C. J. & Wallinga, J. Serial intervals of respiratory infectious diseases: a systematic review and analysis. Am. J. Epidemiol. 180, 865–875 (2014).

68. World Health Organization. Mumps virus nomenclature update: 2012. Weekly Epidemiological Record 87, 217–224 (2012).

69. Müller, N. F., Rasmussen, D. A. & Stadler, T. The Structured Coalescent and Its Approximations. Mol. Biol. Evol. 34, 2970–2981 (2017).

70. Lemey, P., Rambaut, A., Drummond, A. J. & Suchard, M. A. Bayesian phylogeography finds its roots. PLoS Comput. Biol. 5, e1000520 (2009).

71. Gong, L. I., Suchard, M.A. & Bloom, J. D. Stability-mediated epistasis constrains the evolution of an influenza protein. Elife 2, e00631 (2013).

72. Minin, V. N. & Suchard, M. A. Fast, accurate and simulation-free stochastic mapping. Philos. Trans. R. Soc. Lond. B Biol. Sci. 363, 3985–3995 (2008).

73. Bedford, T. Posterior Analysis of Coalescent Trees (PACT).

74. Rota, J. S. et al. Comparison of the Sensitivity of Laboratory Diagnostic Methods from a Well-Characterized Outbreak of Mumps in New York City in 2009. Clin. Vaccine Immunol. 20, 391–396 (2013).

